# An analytical theory of balanced cellular growth

**DOI:** 10.1101/607374

**Authors:** Hugo Dourado, Martin J. Lercher

## Abstract

The biological fitness of unicellular organisms is largely determined by their balanced growth rate, i.e., by the rate with which they replicate their biomass composition. Natural selection on this growth rate occurred under a set of physicochemical constraints, including mass conservation, reaction kinetics, and limits on dry mass per volume; mathematical models that maximize the balanced growth rate while accounting explicitly for these constraints are inevitably nonlinear and have been restricted to small, non-realistic systems. Here, we lay down a general theory of balanced growth states, providing explicit expressions for protein concentrations, fluxes, and the growth rate. These variables are functions of the concentrations of cellular components, for which we calculate marginal fitness costs and benefits that can be related to metabolic control coefficients. At maximal growth rate, the net benefits of all concentrations are equal. Based solely on physicochemical constraints, the growth balance analysis (GBA) framework introduced here unveils fundamental quantitative principles of cellular growth and leads to experimentally testable predictions.

## Introduction

The defining feature of life is self-replication. For non-interacting unicellular organisms in constant environments, the rate of this self-replication is equivalent to their evolutionary fitness^1^: fast-growing cells outcompete those growing more slowly. Accordingly, we expect that natural selection favoring fast growth in specific environments has played an important role in shaping the physiology of many microbial organisms^2, 3^.

Conceptually, we can envision a bacterial cell as a volume enclosed by a membrane, filled with a solution of metabolites and of the proteins and nucleic acids that catalyze their conversion into biomass. A state of the cell is characterized by the molecular concentrations, which in turn determine the fluxes of the biochemical reactions. The boundary conditions limiting the concentrations and fluxes are provided by the environment and by physicochemical constraints. Cellular growth has to be balanced over the cell cycle, i.e., all cellular components must be reproduced in proportion to their abundances^4^. Casting these constraints into a mathematical model and characterizing states of optimal growth may provide a detailed understanding of central aspects of bacterial physiology^3, 5–8^.

Currently, the most popular method to model the physiology of whole cells is flux balance analysis (FBA)^9, 10^. FBA maximizes the production rate of a constant biomass concentration vector while balancing the fluxes producing and consuming internal metabolites to account for mass conservation. All constraints in FBA are linear (Fig. 1). The resulting computational efficiency comes at the price of ignoring reactant concentrations, and hence FBA cannot account for reaction kinetics and the resulting demand for catalytic proteins from first principles. Instead, extensions of FBA that consider enzyme concentrations rely on phenomenological kinetic functions that are assumed to be either constant (FBA with molecular crowding^11^, metabolism and expression models^12^) or linear functions of the growth rate (resource balance analysis^13^).

**Figure 1.**
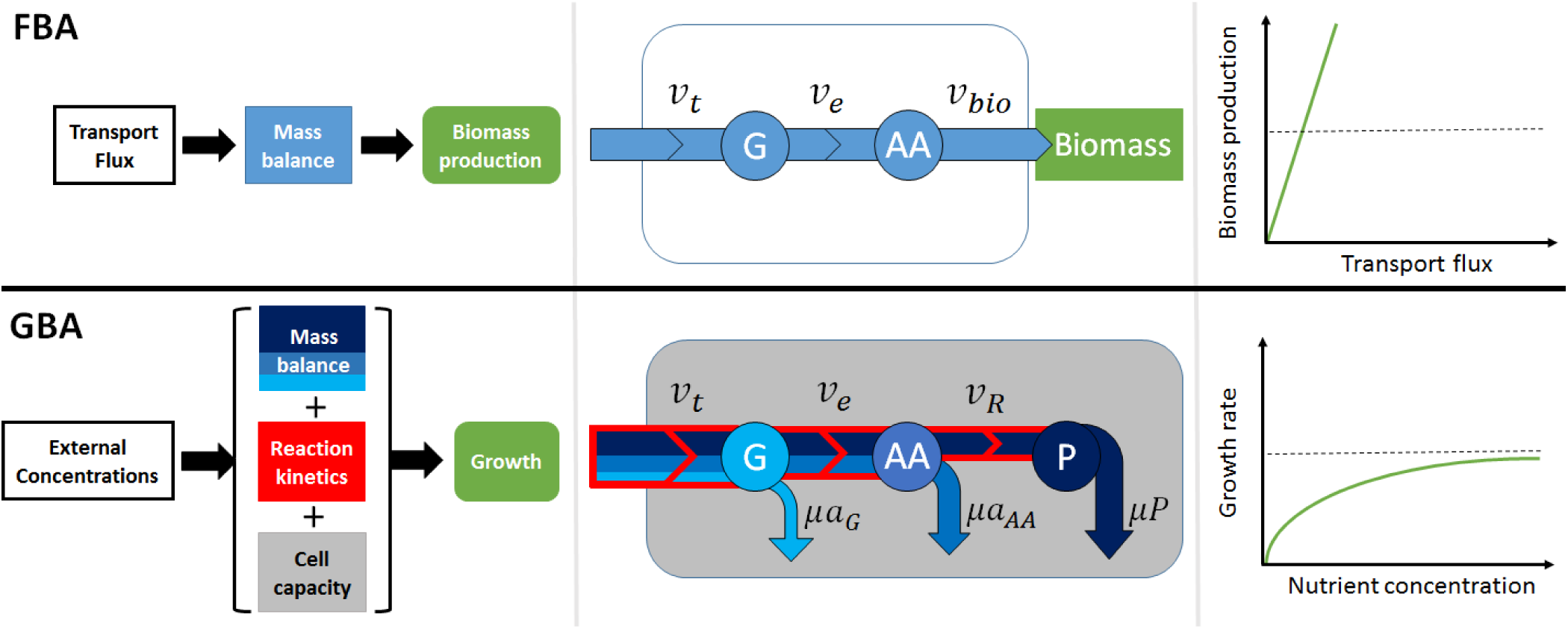
A schematic comparison of flux balance analysis (FBA, top) and growth balance analysis (GBA, bottom) for a simple toy model. A nutrient *G* is taken up through a transporter *t* at rate *v*_*t*_ and is then converted by an enzyme *e* with rate *v*_*e*_ into a precursor for protein synthesis, *AA*. In FBA, *AA* is equated with the biomass, the production of which is maximized while enforcing the stationarity of internal concentrations (blue); this leads to a linear dependence of growth rate on uptake flux. In GBA, *AA* is converted further into total protein *P* by a ribosome *R*, where *P* represents the sum of the three proteins *t, e, R*. GBA maximizes the balanced reproduction of the cellular composition with growth (blue), constrained by non-linear reaction kinetics (red) and cellular capacity (dry mass per volume, grey); this leads to a non-linear dependence of growth rate on nutrient concentrations.

Molenaar et al.^5^ proposed a small, coarse-grained model of balanced growth with explicit non-linear reaction kinetics. Numerical growth rate optimization predicted qualitatively the growth-rate dependencies of cellular ribosome content, cell size, and the emergence of overflow metabolism. No extensions of this approach to models accounting for more than seven cellular reactions have been proposed, likely because of its inherent nonlinearity. Instead, “toy models” of 1-3 reactions were solved analytically to gain further qualitative understanding of systems-level effects^3, 6–8, 14^, including optimal gene regulation strategies^3, 7^.

We term this general modeling scheme *Growth Balance Analysis* (GBA); below, we develop an analytical theory for GBA of arbitrarily complex cellular systems. FBA and its extensions can be viewed as linearizations of the GBA scheme^15^. Fig. 1 shows a schematic comparison of FBA and GBA. While FBA predicts a linear dependence of maximal growth rate on nutrient uptake fluxes, GBA leads to a non-linear (Monod-type) dependence on nutrient concentrations.

## Results

### Modeling balanced exponential growth

Our model assumes that the cell increases exponentially in size, while the concentrations of all cellular components (including the number of membrane constituents per cell volume) remain constant^5^. We do not explicitly model cell division; thus, our model can also be interpreted as describing the growth of a population of cells^7^. In balanced growth, the net production rate of each molecular constituent must balance its dilution by growth, 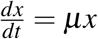, where *x* denotes the concentration of a given component and *µ* is the cellular growth rate^5, 7^.

The mass conservation in chemical reaction networks is commonly described through a *stoichiometric matrix N*, where rows correspond to metabolites and each column describes the mass balance of one reaction^16^. Here, we focus on matrices *A* of active reactions, i.e., *A* is a sub-matrix of *N* that contains all columns *j* for reactions with flux *v* _*j*_ ≠ 0 and all rows for reactants *i* involved in these reactions. *A* also includes a “ribosome” reaction to produce catalytic proteins, encompassing enzymes, transporters, and the ribosome itself. We express concentrations as mass concentrations (mass per volume); accordingly, the entries of *A* are not stoichiometric coefficients but are mass fractions. The mass conservation of each component can then be stated as

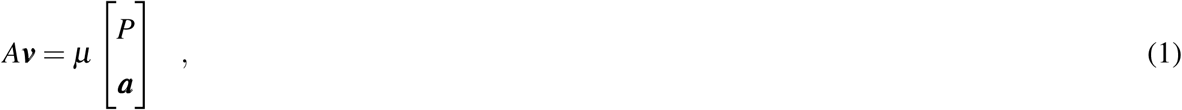

where ***v*** is the flux vector (in units of [mass]/[volume]/[time]), ***a*** is the vector of reactant mass concentrations *a*_*α*_, and *P* is the sum of the mass concentrations *p*_*j*_ of all proteins *j* ∈ {1,…, *n*},

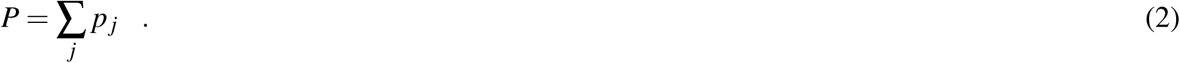

The first row of *A* describes the net production of total protein *P*, which is then distributed among the individual proteins *j*. The remaining rows describe the net production of the reactants *α*.

Each reaction rate *v*_*j*_ is the product of the concentration of its catalyzing protein *p*_*j*_ and a kinetic function *k*_*j*_(***a***) that depends on the reactant concentrations *a*_*α*_,

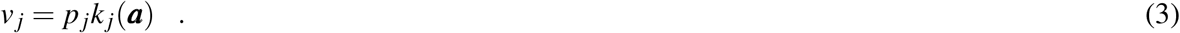

We assume that the functional form and kinetic parameters of *k*_*j*_(***a***) are known. *k*_*j*_(***a***) may depend on the mass concentrations of substrates, products, and other molecules *a*_*α*_ acting as inhibitors or activators, and accounts for the system’s thermodynamics. The activity of all reactions *j* (*v*_*j*_ ≠ 0) implies *p*_*j*_ > 0 and *k*_*j*_(***a***) ≠ 0.

For a given concentration vector ***x*** ≡ [*P*, ***a***]^*T*^, we define a *balanced growth state* (BGS) as a cellular state (characterized by its flux vector ***v***) that satisfies constraints (1), (2), and (3). The set of all such states forms the solution space of balanced growth. On the following pages, we first develop a framework for GBA by characterizing BGSs at a known concentration vector ***x***. Formal definitions and theorems are detailed in **SI text** A.; Table S2 lists the symbols used.

### Cellular state defined by the concentration variables

We define an *elementary growth state* (EGS) as a BGS ***v*** that also represents an elementary flux mode^17^ of a linearized problem (**SI text A.**, Def. 3). We can express any BGS as a weighted average of EGSs at the same concentration vector ***x*** = [*P*, ***a***]^*T*^ (Theorem 3). Moreover, any optimal BGS under a single capacity constraint (see below) is also an EGS (Theorem 9; see also Ref.^18^). Thus, without loss of generality, we focus on EGSs from here on.

If *A* is the active stoichiometric matrix of an EGS, it has full column rank (Theorem 4; see also Ref.^18^). *A* may have more rows than columns, in which case some reactant concentrations are linearly dependent on other concentrations^19^. These dependent concentrations are not free variables, and hence they can be put aside and dealt with separately. For clarity of presentation, we here present only the case without dependent reactants; the generalization can be treated similarly and is detailed in **SI text A.**. Without dependent reactants, *A* is a square matrix with a unique inverse *I* ≡ *A*^−1^. Multiplying both sides of the mass balance constraint (1) by *I*, we obtain (Theorem 5)

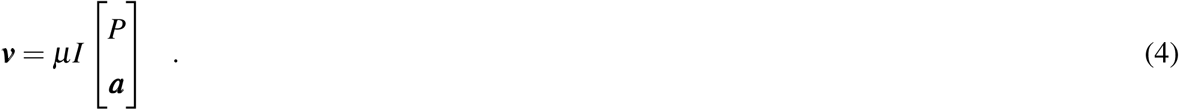

The right hand side of the mass balance constraint (1) quantifies how much of each component needs to be produced to offset the dilution that would otherwise occur through the exponential volume increase. *I*_*ji*_ quantifies the proportion of flux *v*_*j*_ invested into offsetting the dilution of component *i*, and we thus name *I* the *investment* (or dilution) matrix; see Fig. S1 for examples. In contrast to the stoichiometric matrix *A*, which describes local mass balances, *I* describes the structural allocation of reaction fluxes into offsetting the dilution of all downstream cellular components, carrying global, systems-level information.

From the kinetic equations (3), *p*_*j*_ = *v*_*j*_*/k*_*j*_(***a***), and inserting *v*_*j*_ from the investment equation (4) gives

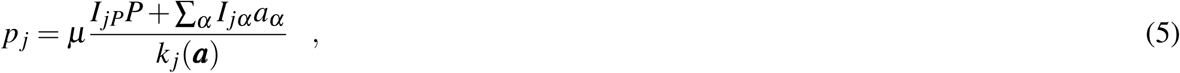

where Σ_*α*_ sums over the set of all reactants (denoted by {*α*}; Theorem 6). Substituting these expressions into the total protein sum (Eq. (2)) and solving for *µ* results in the *growth equation* (Theorem 7)

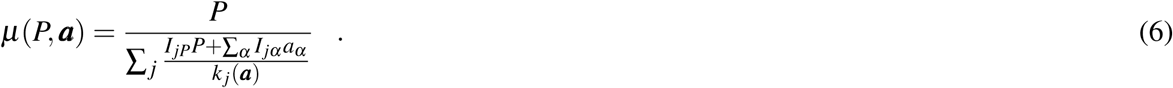

Thus, for any EGS and concentration vector ***x***, there are unique and explicit mathematical solutions for ***v, p***, and *µ*. If *µ* and all individual protein concentrations *p*_*j*_ in Eq. (5) are positive, the cellular state is a BGS; otherwise, no balanced growth is possible at these concentrations.

### Marginal fitness contributions of cellular concentrations

We now use these relationships to calculate the costs and benefits of concentration changes, which are naturally expressed in terms of relative fitness effects. If fitness is determined predominantly by growth rate^1^ (**SI text B.**), we can we define the *marginal net benefit η*_*i*_ of concentration *x*_*i*_ (*i* ∈ {*P, α*}) as the relative change in growth rate^20^ due to a small change in *x*_*i*_ (Def. 4),

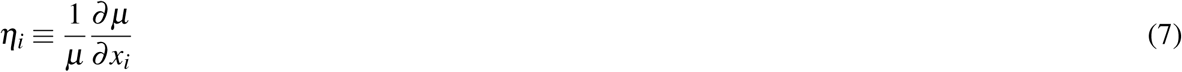

To aid in the interpretation of *η*_*i*_ below, we define the *marginal production cost* incurred by the system via protein *j* as a consequence of increasing concentration *x*_*i*_ at fixed growth rate *µ* and kinetics *k*_*j*_,

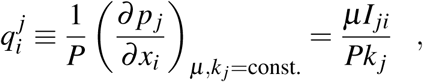

where the second equality arises from the growth equation (6). 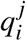 quantifies the proportional increase of *p*_*j*_ to help offset the increased dilution of component *i*. Thus, 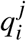 is related to the *protein control coefficient* from metabolic control analysis (MCA); **SI text F.** summarizes the relationship between GBA and MCA^21–23^. From the growth equation (6), it further follows that

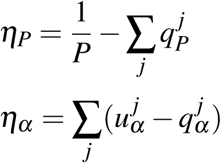

with

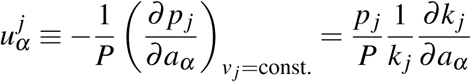

(Theorem 8). 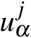 is the *marginal kinetic benefit* of reactant *α* to reaction *j* and quantifies the proportion of protein *p*_*j*_ “saved” due to the change in kinetics associated with an increase in *a*_*α*_ ^24^. *It relates directly to the elasticity coefficients* from MCA (**SI text F.**). The kinetic benefit is nonzero only for reactants that directly affect the kinetics of reaction *j*, making it a purely local effect. Because fluxes are proportional to the concentrations of the catalyzing proteins, the marginal kinetic benefit of total protein is simply *P*^−1^. Overall, the net benefit of component *i* via reaction *j* is the reduction of the protein fraction *p*_*j*_*/P* at constant *µ* facilitated by the increase in *x*_*i*_,

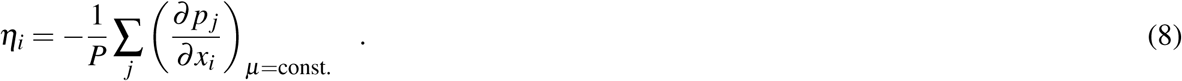

This result provides a formal justification for the notion that cellular costs lie predominantly in protein production^3, 5–8, 12–14, 24–27^.

### Optimal growth and the balance of marginal net benefits

Up to this point, we kept ***x*** = [*P*, ***a***]^*T*^ fixed. We will now characterize optimal growth states, i.e., BGSs with maximal growth rate across all allowed concentration vectors. To make this problem well defined, we need to consider an additional constraint that reflects the cellular requirement for a minimal amount of free water to facilitate diffusion^28, 29^. We implement this constraint by assuming that cellular dry weight per volume is limited to a maximal density *ρ*, where *ρ* is determined by external osmolarity^29, 30^ but is otherwise constant across growth conditions^31^,

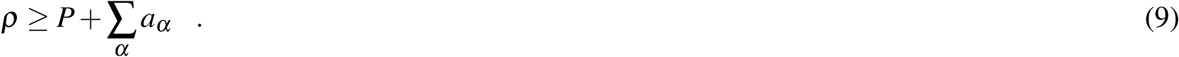

A BGS is a *capacity-constrained balanced growth state* (cBGS) if it additionally satisfies constraint (9). At maximal growth rate, the cellular components will utilize the full cellular capacity to saturate enzymes with their substrates, and thus the inequality in Eq. (9) becomes an equality.

The maximal obtainable balanced growth rate *µ*\* will be a function of *ρ*. In analogy to the marginal net benefits of cellular components, we define the *marginal benefit* of the cellular capacity as the fitness increase facilitated by a small increase in *ρ*,

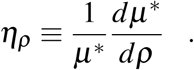

Using the method of Lagrange multipliers with the growth equation (6) as the objective function, we derive necessary conditions at optimal growth, which we term *balance equations*:

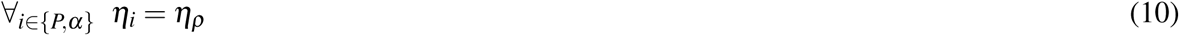

(Theorem 10). The optimal state is perfectly balanced: the marginal net benefits of all cellular concentrations *x*_*i*_ are identical. If the dry weight density *ρ* could increase by a small amount (such as 1 mg/l), then the marginal fitness gain that could be achieved by increasing protein concentration by this amount is identical to that achieved by increasing the concentration of any reactant *α* by the same amount. This should not be surprising: if the marginal net benefit of concentration *x*_*i*_ was higher than that of *x*_*i*′_, growth could be accelerated by increasing *x*_*i*_ at the expense of *x*_*i*′_.

Eq. (10) together with Eq. (9) describes a system of *n* + 1 equations for *n* + 1 unknowns. In realistic cellular systems, this set of equations has a finite number of discrete solutions. Thus, growth rate optimization can be replaced by searching for the solution of the balance equations. If the optimization problem is convex, the conditions given by Eq. (10) are necessary and sufficient, and the solution is unique.

### Quantitative predictions

If a substrate *i* is consumed only by a single reaction that produces *i*′, the non-local dilution terms in the balance equation *η*_*i*_ = *η*_*i*′_ cancel, and we are left with a local problem for which only the kinetic benefits of *x*_*i*_ and *x*_*i*′_ must be considered. This is the case for protein production in simplified models where the ribosome (*R*) produces proteins from a single substrate, a generic ternary complex (*T*)^26^. In such models, we can calculate the optimal protein fraction of actively translating ribosomes, *ϕ*_*R*_ = *p*_*R*_*/P* from the balance equation *η*_*T*_ = *η*_*P*_ (**SI text C.**), which agrees quantitatively with experimental values^32, 33^ (Fig. 2).

**Figure 2.**
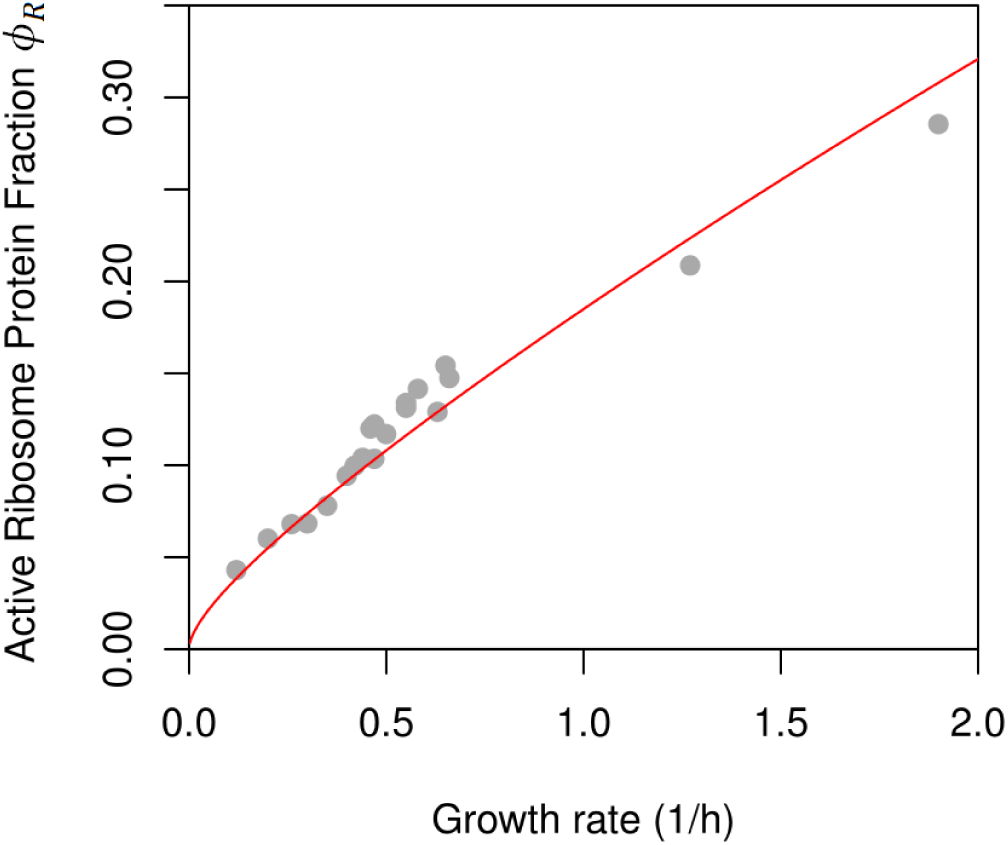
The GBA prediction of the active ribosomal protein fraction in *E. coli* agrees with experimental values. Comparison of GBA predictions (red line, no free parameters) and data across 20 different growth conditions^32, 33^ (grey dots) results in a Pearson correlation coefficient *R*^2^ = 0.96 (*P <* 10^−13^).

An approximation that ignores the dilution of intermediates and hence production costs 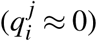 results in less accurate predictions especially at high growth rates (Fig. S4). In growth on minimal media (*µ <* 1), the dilution of intermediates *µa*_*α*_ becomes less important. This explains why the relationship between the concentrations of a substrate and its catalysts is well approximated in this regime through minimizing their joint utilization of cellular capacity^34^.

To obtain a rough quantitative estimate of the marginal net benefits *η*_*i*_, we here consider the simplest model of a complete cell, consisting of only a transport protein and the ribosome^3, 6^ (Fig. S2). Based on the experimentally observed protein fraction of total dry weight in *E. coli, P/ρ* = 0.54,^29^ we estimate 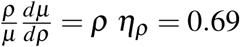 (**SI text D.**). Thus, a decrease in cellular dry weight density *ρ* of 1% would lead to a 0.69% decrease in growth rate, emphasizing the biological significance of the capacity constraint.

The cellular capacity *ρ* changes when external osmolarity is modified^29^. 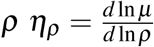 is the slope of the log-scale plot of *µ* vs. *ρ* at different external osmolarities. While increases in *ρ* may have strong effects on diffusion and thus on enzyme kinetics, reductions in *ρ* due to decreased external osmolarity are within the scope of our model. The very limited available experimental data (three data points from Ref.^35^, Fig. S3) suggests *ρ η*_*ρ*_ ≈ 0.66, close to our rough estimate from the minimal cell model.

## Discussion

Our derivations are based on the insight that for any EGS, the inverse *I* of the active stoichiometric matrix quantifies how individual fluxes offset the dilution of downstream cellular components by growth. These non-local, structural constraints arising from mass balance lead to an explicit dependence of reaction fluxes on the cellular concentrations (Eq. (4), Theorem 5). Independently of this, fluxes also depend on concentrations through reaction kinetics (constraint (3)). Combining these two relationships leads to explicit expressions for the individual protein concentrations *p*_*j*_ and for the growth rate *µ* as functions of arbitrary concentrations [*P*, ***b***]^*T*^. As any BGS can be expressed as a weighted average of EGSs (Theorem 3), these results allow a general characterization of the solution space of balanced growth. Further, the growth rate equation (6) can be employed to calculate marginal fitness benefits of concentrations and to derive balance equations for marginal benefits at optimal growth.

Previous work has emphasized the central role of proteins in the cellular economy^3, 5–8, 12–14, 24–27^, and this is confirmed by Eq. (8). However, whereas total protein mass concentration in real biological systems is indeed much higher than the mass concentration of any other cellular constituent *a*_*α*_, the balance equations show that at optimal growth, their marginal net benefits are in fact equal.

To make the presentation concise, our development of GBA assumes (i) that all proteins contribute to growth by acting as catalysts or transporters; (ii) that there is a 1-to-1 correspondence between proteins and reactions; (iii) that proteins are not used as reactants; (iv) that all catalysts are proteins; and (v) that cells are optimized for growth. As outlined in **SI text E.**, it is straight-forward to remove these simplifications.

In principle, exploitation of the balance equations (Eq. (10)) may allow the numerical optimization of cellular systems of realistic size, encompassing hundreds of protein and reactant species. One remaining obstacle to the accurate formulation of such models is the current incompleteness of the kinetic constants needed to parameterize the functions *k*_*j*_**(*a***). Until methods for the high-throughput ascertainment of kinetic parameters^36^ are fully developed, artificial intelligence may provide reasonable approximations^37^ for the required parameters. As an alternative to genome-scale models, the balanced growth theory developed here could be applied to coarse-grained cellular models of increasing complexity, parameterized from experimental data^27, 38^.

Our work extends previous ad-hoc optimizations of toy models^3, 5–8^ into a theory of balanced growth. We show that the balanced growth framework allows general, quantitative insights into cellular resource allocation and physiology, as exemplified by the growth and balance equations. Application and further development of this theory may foster an enhanced theoretical understanding of how physicochemical constraints determine the fitness costs and benefits of cellular organization. Moreover, the explicit expressions for the (marginal) costs and benefits of cellular concentrations in terms of fitness provide a rigorous framework for analyzing the cellular economy. We anticipate that this approach will prove fruitful in the interpretation of natural and laboratory evolution, and in optimizing the design of synthetic biological systems.

## Acknowledgements

We thank Johannes Berg, Oliver Ebenhöh, Daan de Groot, Xiao-Pan Hu, Terry Hwa, Michael Lässig, Wolfram Liebermeister, Elad Noor, and Deniz Sezer for discussions. This work was funded by the Deutsche Forschungsgemeinschaft (DFG, German Research Foundation) through grants IRTG 1525, CRC 680, CRC 1310, and, under Germany’s Excellence Strategy, through grant EXC 2048/1 (Project ID: 390686111).

## Author contributions

HD and MJL jointly conceived the study, interpreted the results, and wrote the manuscript. HD developed the GBA framework, performed all data analyses, and derived all formal results except Theorems 1-3, which were derived by MJL.

## Supplementary Information

### A. Growth balance analysis

In this section, we provide a formal description of growth balance analysis (GBA), detailing the formal definitions, theorems, and proofs that form the basis of the main text. For simplicity of notation, we use the following conventions: {*α*} is the set of all reactants in the active stoichiometric matrix *A*, and Σ_*α*_ indicates that we sum over all *α* ∈ {*α*}. We use corresponding notations for the sets of *basis reactants* {*β*}, with concentrations *b*_*β*_, and *dependent reactants* {*γ*}, with concentrations *c*_*γ*_ (see below).

#### Characterization of balanced growth states

First, we introduce the fundamental definitions that characterize the solution space of balanced cellular growth. We define balanced growth states and generalize the concept of elementary flux modes from linear constraint-based models to elementary growth states (defined as flux vectors). We then introduce several theorems on the characterization and decomposition of balanced growth states.

In the formulation presented here, we assume that proteins do only act as catalysts and not as substrates of reactions. Hence, neither total protein nor individual proteins are considered “reactants”.

##### Definition 1

(Balanced growth states (BGSs)). *Let* ***v*′** ∈ ℝ^*n*′^ *be the vector of fluxes through the biochemical reactions that occur in a cell, in units of [mass/volume/time]. Let 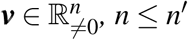, be the subvector of* ***v***′ *that contains all active fluxes of* ***v***′ *(i.e., all entries 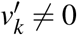)*. *Let 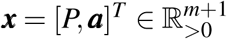 be a corresponding vector of total protein concentration P and individual reactant concentrations a*_*α*_, *α* ∈ {1,…, *m*}., *where each a*_*α*_ *is consumed or produced by at least one of the fluxes v*_*i*_; *x*_*i*_ *is in units of [mass/volume]. Let A* ∈ ℝ^(*m*+1)×*n*^ *be the corresponding active stoichiometric matrix in mass fraction units, i.e., column j of A describes reaction j with flux v*_*j*_, *row i of A corresponds to the cellular component x*_*i*_, *and each column is mass balanced. Thus, the sum of negative entries in each column is S*_−_ = −1 *and the sum of positive entries of each column is S*_+_ = +1; *for reactions that involve an external substrate not represented by a row of A*, −1 *< S*_−_ ≤ 0, *while for reactions that involve an external product*, 0 ≤ *S*_+_ *<* 1.

*Let 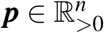 be the vector of individual protein concentrations (in units of [mass/volume]), where protein j catalyzes reaction j; for simplicity, we assume that “ribosome” catalyzing protein production is also itself a protein. Let* ***k***(***a***) *be a vector of kinetic functions, 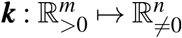, where k*_*j*_(***a***) *is in units of [1/ time].*

*Then* ***v*** *is a* ***balanced growth state (BGS)*** *at growth rate µ if and only if it fulfills the following three constraints:*

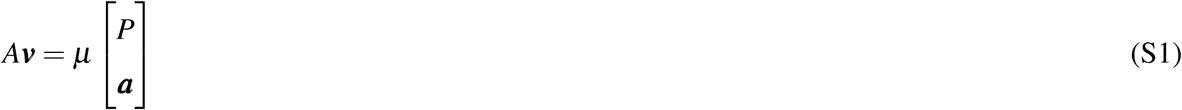

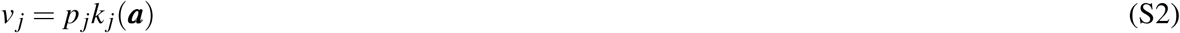

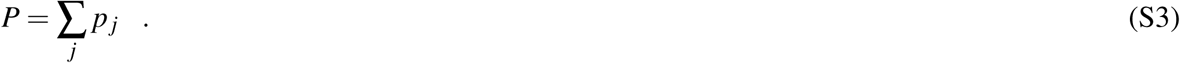

*Constraint (S1) implements mass balance, constraint (S2) implements concentration-dependent reaction kinetics, while constraint (S3) implements a constraint on the total proteome concentration.*

*A BGS* ***v*** *at growth rate µ is a* ***capacity-constrained balanced growth state (cBGS)*** *if it additionally fulfills the constraint on cellular capacity*

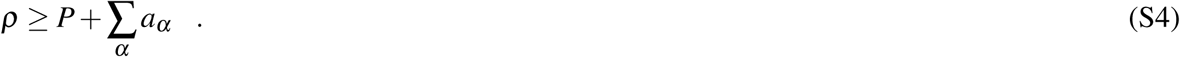

The kinetic constraint (S2) assumes that the flux through each reaction is linear in the concentration of the catalyzing enzyme, while the dependence on the reactant concentrations *a*_*α*_ will typically be non-linear. For simplicity of notation, we will sometimes make the dependence of kinetics on ***a*** implicit, i.e., we will use *k*_*j*_ ≡ *k*_*j*_(***a***).

In the above definitions, we define BGSs (or cBGSs) as a function of the set of active reactions (corresponding to the columns of *A*) and the concentration vector. The set of all such states at all concentrations 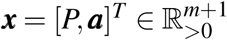 defines the solution space of balanced growth (or capacity-constrained balanced growth) for a given active stoichiometric matrix *A*.

If we consider the concentration vector ***x*** = [*P*, ***a***]^*T*^ as a descriptor of a constant biomass composition, Eq. (1) is mathematically identical to the flux balance equation at the heart of FBA (see., e.g., Ref.^39^).

Based on biophysical considerations, we might replace Eq. (S4) with separate capacity constraints on the total volume concentration inside each cellular compartment^28^ and on the total area occupied by non-lipid membrane components per membrane area^5, 40^. An even simpler capacity constraint imposed in most previous models^3, 5–8, 12–14^ is to fix total protein concentration *P* to a constant value. However, it has been shown that *P* decreases with increasing growth rate^31, 41^. Thus, while a constant *P* allows to simplify the presentation, Eq. (9) provides a more meaningful constraint; moreover, Eq. (9) allows us to determine the costs and benefits of varying the total protein concentration.

De Groot *et al.* have defined balanced growth states for a similar problem^18^. In their formulation, the dimensions of the concentration vector ***x*** include not only total protein *P*, but all individual protein concentrations *p*_*j*_. This more general problem formulation comes at the cost of more involved decomposition rules^18^ compared to Theorem 2, and does not lend itself to the derivation of explicit expressions for growth rate (Theorem 7), fitness costs of concentrations (Theorem 8), or necessary conditions of maximal balanced growth (Theorem 10).

We now provide the basis for linking BGSs to elementary flux modes, which are defined for FBA-type linear constraint-based problems^17^ and which have been extended to proteome-constrained models^42, 43^.

##### Definition 2

(Elementary flux modes (EFM)). 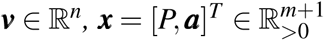, *and A* ∈ ℝ^(*m*+1)×*n*^ *be as in Def. 1. Let 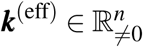 be a vector of effective kinetic constants. Then we call* ***v*** *a* ***feasible flux vector*** *at biomass production rate v*_*bio*_ *if and only if it fulfills the following constraints:*

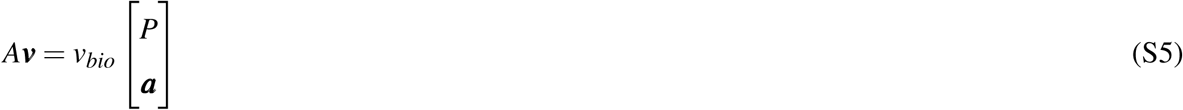

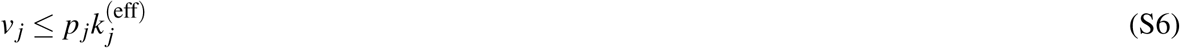

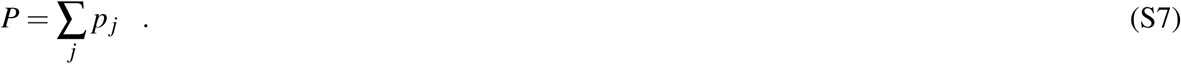

*A feasible flux vector* ***v*** *is a representative of an* ***elementary flux mode*** *if and only if it is non-decomposable, i.e., it fulfills the following additional constraint*^*17*^: *There exists no couple of feasible flux vectors* ***v***′, ***v***′′ *such that* ***v*** = *λ*_1_ ***v***′ + *λ*_2_ ***v***′′ *with λ*_1_, *λ*_2_ > 0 *and where both* ***v***′ *and* ***v***′′ *have at least the same number of zeroes as v, while at least one of them contains more zeroes than v.*

Constraint (S5) is equivalent to the standard steady state constraint of flux balance analysis problems, formulated with an equation analogous to Eq. (S1) for a fixed biomass vector instead of including an artificial “biomass reaction” in *A* (see, for example, Eq. (2) in Ref.^39^).

Note that in the definition of EFMs, both the biomass composition ***x*** = [*P*, ***a***]^*T*^ and the effective kinetics ***k***^(eff)^ are assumed to be constant; thus, the constraints (S5)-(S7) that define the space of feasible flux vectors are fully linear. In contrast, constraint (S2) in Def. 1 defines reaction kinetics as a function of the reactant concentrations ***a***.

##### Definition 3

(Elementary growth state (EGS)). *A BGS* ***v*** *is an* ***elementary growth state*** *(EGS) if and only if it is a representative of a corresponding EFM, i.e.*, ***v*** *represents an EFM of the corresponding linear problem with constant biomass* ***x*** = [*P*, ***a***]^*T*^ *and effective kinetic constants* ***k***^(eff)^ = ***k***(***a***).

We emphasize that ***v*** is an EFM of the corresponding linearized (FBA-like) problem (see Def. 2), not of the balanced growth problem from Def. 1 from which it is derived. EFMs are defined as equivalence classes of minimal feasible steady-state flux distributions, whose members can be converted into each other by multiplication with a positive scalar^17^. This definition cannot be generalized to balanced growth models, as multiples of an admissible flux vector generally do not satisfy constraint (S1). For this reason, de Groot *et al.* have generalized the concept of EFMs to equivalence classes of minimal sets of active reactions in balanced growth states, termed *elementary growth modes* (EGMs)^18^.

##### Theorem 1

(Existence of solutions). *Let* ***x*** = [*P*, ***a***]^*T*^ *be a concentration vector, and µ* > 0 *be a growth rate. For any flux vector* ***v***′ *that satisfies the mass balance constraint (S1), there exists a unique BGS* ***v*** = *λ* ***v***′ *with λ* > 0 *if all fluxes run in the direction compatible with the reaction kinetics (i.e.*, ∀_*j*_ *k*_*j*_*v*_*j*_ > 0*), and no such BGS otherwise.*

*Proof.* From constraint (S2), it is clear that if *k*_*j*_*v*_*j*_ ≤ 0, no BGS with *p*_*j*_ > 0 exists. For *k*_*j*_ ≠ 0, the concentration of protein *j* is uniquely defined by *p*_*j*_ = *v*_*j*_*/k*_*j*_ (constraint (S2)). Let *P*′ = Σ _*j*_ *v*′*j/k*_*j*_ be the total protein concentration associated with ***v*′**. Then setting *λ* ≡ *P/P*′ results in the only flux vector that fulfills all constraints of Def. (1).

Next, we use this result to show that any weighted average of BGSs is itself a BGS.

##### Theorem 2

(A weighted average of BGSs is a BGS). *Let* (***v***^(1)^, …, ***v***^(*k*)^) *be an ordered set of BGSs for the concentration vector* ***x*** = [*P*, ***a***]^*T*^ *with growth rates* (*µ*^(1)^,…, *µ*^(*k*)^), *but with potentially different active stoichiometric matrices A*^(*l*)^. *Let A be the stoichiometric matrix that combines all reactions represented in* (*A*^(1)^, …, *A*^(*k*)^), *i.e., the columns of A consist of all unique columns of* (*A*^(1)^, …, *A*^(*k*)^). *Let* (***v***′^(1)^, …, ***v***′^(*k*)^) *be a representation of the individual BGSs* ***v***^(*l*)^ *in the flux space defined by A, i.e.*, 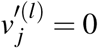 *for all columns (reactions) of A not represented in A*^(*l*)^. *Then any weighted average* ***v*** = Σ_*l*_ *w*_*l*_ ***v***′^(*l*)^ *of these extended flux vectors (with weights w*_*l*_ > 0 *and* Σ_*l*_ *w*_*l*_ = 1*) is itself a BGS for* ***x***, *with a growth rate that is the weighted average of the individual growth rates, µ* = Σ_*l*_ *w*_*l*_ *µ*^(*l*).^

*Proof.* The mass balance constraint (S1) is linear in the fluxes and growth rates, and is hence also fulfilled for the weighted averages. The protein concentrations of each BGS ***v***′^(*l*)^ are 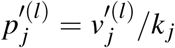. To satisfy the reaction kinetics constraint (S2), the protein concentrations of the weighted average are 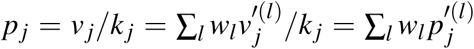. As each BGS (*l*) fulfilled the proteome constraint (S3), *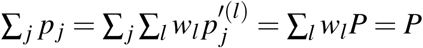*, and thus ***v*** is a BGS.

We can now use Theorems 1 and 2 together with results on elementary flux modes to show that any BGS can be decomposed into a weighted average of EGSs.

##### Theorem 3

(BGSs are weighted averages of EGSs). *Any BGS* ***v*** *for the concentration vector* ***x*** = [*P*, ***a***]^*T*^ *can be decomposed into a weighted average of EGSs at x.*

*Proof.* ***v*** is a feasible flux vector for the linearized problem defined by constraints (S5)-(S7) at constant biomass ***x***. The direction of reaction *j* is fixed by the sign of 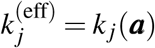, i.e., all reactions are irreversible. Under these conditions, it has been shown that ***v*** is a convex combination of elementary flux modes ***v***′^(*l*)^ of the linear problem^17^, i.e., 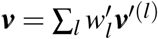 with 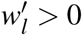. From Theorem 1, we know that for each of these EFMs, there exists a unique BGS ***v***^(*l*)^ = *λ*_*l*_ ***v***′^(*l*)^ with *λ*_*l*_ > 0; according to Def. 3, this is an EGS. Thus, we can write ***v*** = Σ_*l*_ *w*_*l*_ ***v***^(*l*)^ as a linear combination of EGSs, with weights 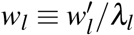.

To prove that ***v*** is a weighted average of the ***v***^(*l*)^, it remains to be shown that *W* ≡ Σ_*l*_ *w*_*l*_ = 1. According to Theorem 2, a weighted average 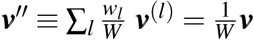 will also be a BGS. However, Theorem 1 states that there exists only one BGS in the direction of ***v***, and thus *W* = 1.

#### Growth equations

In this section, we assume that the concentrations of total protein and of individual reactants, ***x*** ≡ [*P*, ***a***] are known. Mass conservation (constraint (S1)) and reaction kinetics (constraint (S2)) relate reaction fluxes to the concentration vector in two fundamentally different ways. We will now exploit this fact to eliminate the flux variables and to derive explicit expressions for ***v, p***, and *µ*.

Note that because the concentrations ***x*** are used as input parameters in these analyses, no explicit consideration of constraints on cellular capacity, such as constraint (S4) is necessary. The given concentrations ***x*** may obey constraint (S1) or alternative capacity constraints, such as independent constraints on the capacity of cellular compartments, but these will not be used here. They will only become important when we vary ***x*** to find states of maximal growth rate in Section A..

An important requirement for the analyses below is that the active stoichiometric matrix *A* has full column rank, motivating the next theorem.

##### Theorem 4

(The active reactions of an EGS are linearly independent). *Let A* ∈ ℝ^(*m*+1)×*n*^ *be the active stoichiometric matrix of an EGS. Then A has full column rank n, i.e., the columns of A are linearly independent.*

*Proof.* According to the definition of EGSs (Def.3), *A* is also the active matrix of the corresponding linearized (flux balance type) problem. It has previously been shown that the active stoichiometric matrix *A* of an EFM of a linear flux-balance problem has full column rank if *A* is formulated without an explicit “biomass” reaction (as in Def. 2)^44^.

According to this theorem, the following theorems - which assume that *A* has full column rank - can in particular be applied to EGSs (and, as we will see below in Theorem 9, thus also to cBGSs with maximal growth rate).

##### Theorem 5

(Investment equation). *Let A* ∈ ℝ^(*m*+1)×*n*^ *be an active stoichiometric matrix of a flux vector* ***v*** *that fulfills the mass balance constraint (S1) with concentration vector* ***x*** = [*P*, ***a***]^*T*^, *where A has full column rank n. Then we can split A into two submatrices B* ∈ ℝ^*n*×*n*^ *and C* ∈ ℝ^(*m*+1−*n*)×*n*^,

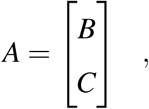

*such that B is a non-singular (invertible) square matrix and each row of C is a linear combination of rows of B. Let I* ≡ *B*^−1^. *Let* ***b*** *be the subvector of reactant concentrations* ***a*** *that correspond to the rows of B, and let* ***c*** *be the subvector of the remaining reactant concentrations. Then* ***v*** *is given by*

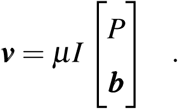

*The dependent reactant concentrations* ***c*** *are linear combinations of the independent concentrations* [*P*, ***b***]^*T*^,

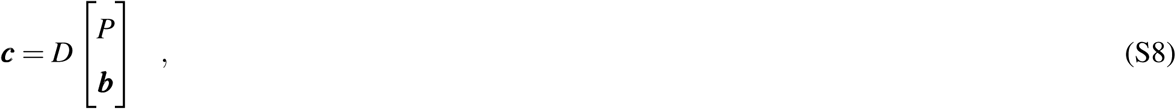

*with the dependence matrix D* ≡ *CI.*

*Proof. A* may have more rows than columns (*m* + 1 > *n*). In this case, the rows for exactly *n* metabolites are linearly independent, as row and column rank must equal. As a consequence, the remaining *m* + 1 −*n* metabolite concentrations are linearly dependent on the concentrations of the *n* independent metabolites. These dependent concentrations are not free variables, and hence they can be put aside and dealt with separately.

We decompose the linear system of equations represented by constraint (S1) into two parts, rearranging the rows of *A* into matrices *B,C* such that *B* contains the rows for the independent reactants. As *A* has full column rank, choosing linearly independent rows results in a square matrix *B* of full rank (#*rows*(*B*) = *rank*(*B*) = *rank*(*A*)). Let ***b*** be the subvector of reactant concentrations ***a*** that correspond to the rows of *B*, and let ***c*** be the subvector of the remaining reactant concentrations corresponding to the rows of *C*. We can then split the mass balance constraint (S1) into two separate equations:

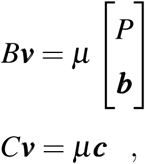

*B* is a square matrix of full rank, so there is always a unique inverse *I* ≡ *B*^−1^. Multiplying both sides of the first equation by *I* from the left, we obtain the desired equation for ***v***. Inserting this result into the second equation results in the desired equation for ***c***.

Thus, if *A* has full rank, then any flux vector ***v*** respecting the flux balance constraint (S1) is uniquely defined and is a linear combination of the total protein concentration *P* and the independent metabolite concentrations ***b***. Each entry of the inverse matrix *I*_*ji*_ quantifies the proportion of flux *j* invested into the dilution of component *i*, and we thus name *I* the *investment* (or *dilution*) matrix (see Fig. S1 for examples). In contrast to the stoichiometric matrix *A*, which describes local mass balances (constraint (S1)), *I* describes the structural allocation of reaction fluxes into the production of cellular components diluted by growth, and thus carries global, systems-level information.

*B* corresponds to the reduced stoichiometric matrix in Ref.^19^. *D* describes the linear dependence of the *dependent concentrations* ***c*** on *P* and ***b***; it is identical to the link matrix in Ref.^19^. The relationship between *A* and *B,C* can be understood in terms of matroid theory, where the rows of *B* form a *basis* for the *matroid* spanned by the rows of *A*, and the set of rows of *C* is the *closure* for the set of rows of *B*. If the choice for the partitioning of *A* into *B* and *C* is not unique, some partitionings may be pathological and should be avoided (**SI text G.**).

When *A* is not square, *B* includes a proper subset of the rows in *A*, and thus *B* on its own is not mass balanced. The “missing” mass fluxes are balancing ***c***, and hence the flux investment into ***c*** is already accounted for by by the investment equation in Theorem 5.

We are now in a position to express the individual protein concentrations and the growth rate of a BGS as explicit functions of the concentrations ***x*** = [*P*, ***a***]^*T*^.

##### Theorem 6

(Individual protein concentrations as a function of the independent concentrations). *Let A* ∈ ℝ^(*m*+1)×*n*^ *be an active stoichiometric matrix with full column rank n, and let* ***x*** = [*P*, ***a***]^*T*^ *be a concentration vector. Let* ***v*** *be a corresponding BGS. Let B and D be the basis and dependency matrices, respectively, as defined in Theorem 5, and let I* = *B*^−1^. *Then the concentration of the protein catalyzing reaction j is*

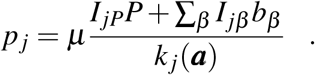

*Proof.* As *A* is an active matrix, all fluxes *v*_*j*_ = *p*_*j*_*k*_*j*_(***a***) (constraint (S2)) are non-zero. We can thus express the individual protein concentrations as *p*_*j*_ = *v*_*j*_*/k*_*j*_(***a***). Inserting *v*_*j*_ from the investment equation (Theorem 5) directly leads to the above equation.

We now insert the equations for the individual proteins into the total protein constraint (S3) to obtain an explicit expression for the growth rate.

##### Theorem 7

(Growth equation). *Let A* ∈ ℝ^(*m*+1)×*n*^ *be an active stoichiometric matrix with full column rank n, and let* ***x*** = [*P*, ***a***]^*T*^ *be a concentration vector. Let* ***v*** *be a corresponding BGS. Let B and D be the basis and dependency matrices, respectively, as defined in Theorem 5, and let I* = *B*^−1^. *Then the growth rate is*

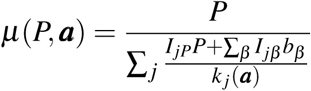

*if for all reactions* 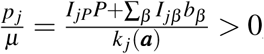, *and no balanced growth is possible otherwise.*

*Proof.* According to Theorem 6, the individual protein concentrations are 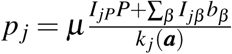. The flux *v*_*j*_ catalyzed by protein *j* must be active, and thus *p*_*j*_ has to be positive for all *j*. Substituting the expressions for *p*_*j*_ into the proteome constraint (S3), we obtain

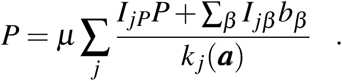

The sum on the r.h.s. is positive, and dividing by it results in the growth equation.

Thus, if the active matrix *A* of a BGS is full rank, there are unique and explicit mathematical solutions for ***p, v***, and *µ*. In particular, this is the case for optimal growth states, as well as for all other EGSs. In this section, we did not impose any capacity constraints (such as constraint (S4)), and thus Theorems 1-7 remain valid under arbitrary capacity constraints (as long as the capacity constraints are respected by the concentration vector ***x*** = [*P*, ***a***]^*T*^).

#### Marginal fitness benefits and costs

In this section, we first define marginal fitness benefits and costs of concentrations. As in the previous section, the definitions make no use of the capacity constraint (S4), and thus remain valid under alternative capacity constraints. After introducing the definitions, we explore the marginal fitness benefits of cellular concentrations at optimal growth; at this point, the capacity constraint becomes central to our analysis.

##### Definition 4

(Marginal costs and benefits). *Let* ***v*** *be a BGS with growth rate µ. Let i* ∈ {*P, β*} *be an index of the concentration vector* ***x*** = [*P*, ***b***]^*T*^.

*Then the* ***direct marginal net benefit*** *of concentration x*_*i*_ *is defined as the relative change in growth rate due to a small change in x*_*i*_^*20*^,

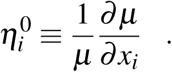

*The* ***marginal production cost*** *of x*_*i*_ *is defined as*

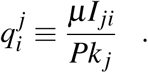

*The* ***marginal kinetic benefit*** *of x*_*i*_ *is defined as*

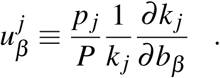

*Further, we define the (total) marginal kinetic benefit of dependent reactant γ as*

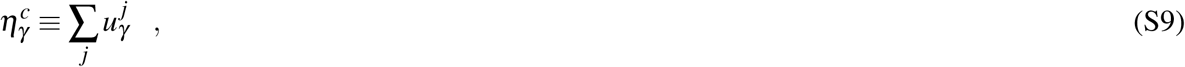

*with the* ***marginal kinetic benefit*** *of reactant γ to reaction j*

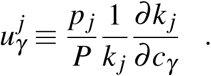

*The (total)* ***marginal net benefit*** *of x*_*i*_ *is then defined as the relative change in growth rate due to a small change in x*_*i*_, *accounting for the resulting changes in the concentration of dependent metabolites c*_*γ*_,

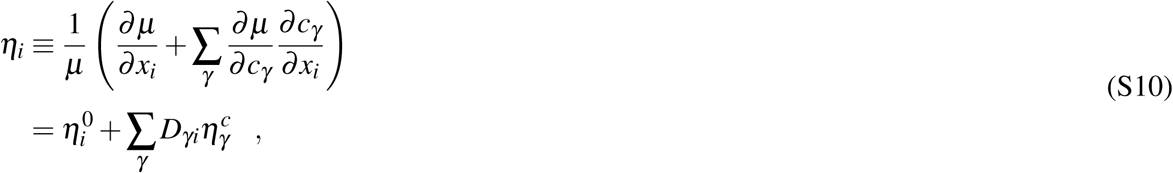

*where the second equality follows from Eq. (S8), Theorem 7, and the previous definitions.*

*A change δx*_*i*_ *of x*_*i*_ *(i* ∈ {*P, β*}*) causes a correlated change of each dependent concentration δc*_*γ*_ = *D*_*γi*_*δx*_*i*_ *(Eq. (S8)). Thus, a change by δx*_*i*_ *results in a total change of the utilization of cellular capacity by κ*_*i*_*δx*_*i*_, *with the* ***capacity factor*** *defined as*

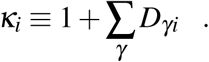

The definition of 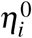 accounts for the production costs of dependent reactants *c*_*γ*_, as these costs are embedded in *I* (*B* is not balanced if there are dependent reactants). However, 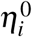 ignores the kinetic benefits of the dependent reactants; this is why the definition of *η*_*i*_ includes a separate term for their kinetic benefits but not their costs.

If *I*_*ji*_ and *k*_*j*_ are both positive, then the production cost 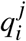 is also positive, i.e., it decreases fitness. The production costs are global, systems-level effects, quantified through the investment matrix *I*. In contrast, the kinetic benefit 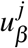 is a local effect, as it is non-zero only for reactants *β* directly involved in reaction *j*. 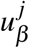 will generally be positive if *β* is a substrate of reaction *j*.

The marginal net benefits can be expressed as the difference between marginal benefits and costs.

##### Theorem 8

(Direct marginal net benefits). *The direct marginal net benefits of the total protein concentration P and of independent reactant concentrations b*_*β*_ *(β* ∈ {1,…, *m*}*), respectively, are*

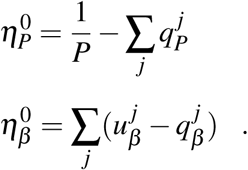

*Proof.* Taking the corresponding derivatives (see Def. (4)) of the growth equation (Theorem 7) directly leads to these equations.

So far, we have considered BGS for a given set of active reactions (corresponding to the columns of *A*) and given concentrations ***x*** = [*P*, ***a***]^*T*^. Below, we will examine capacity-constrainted BGSs (cBGSs) with maximal growth rate given the set of active reactions, optimized over all concentration vectors 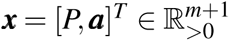 that respect the capacity constraint (S4). As a preparation for these analyses, we first show that cBGSs at optimal growth are EGSs.

##### Theorem 9

(cBGSs with maximal growth rate are EGSs). *Let N be a stoichiometric matrix of a general balanced growth model. Let* ***v***\* *be a cBGS that maximizes the growth rate of the general problem. Then* ***v***\* *is an EGS.*

*Proof.* Without loss of generality, we restrict ***v***\* to its active dimensions 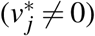, with active stoichiometric matrix *A*. Then this reduced ***v***\* is the optimal solution for the following non-linear optimization problem over all concentration vectors 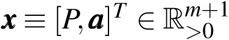:

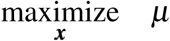

subject to:

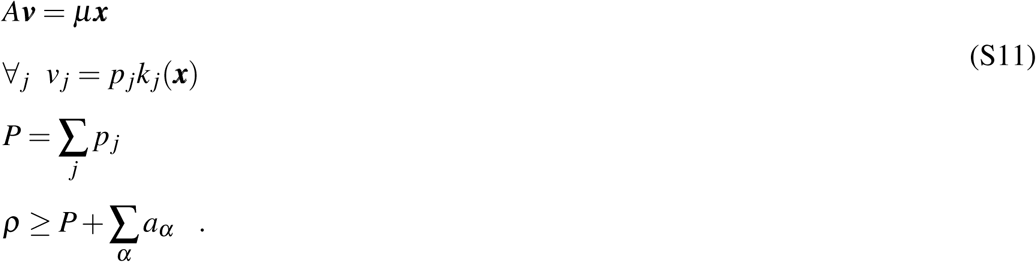

Let ***x***\* = [*P*\*, ***a***\*]^*T*^ be the concentrations and *µ*\* the growth rate of the optimal solution ***v***\*. Now let us consider a linearized version of this optimization problem, where me maximize the production rate *v*_*bio*_ at constant biomass composition ***x***\* and effective kinetic constants 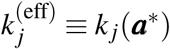 (see Def. 2):

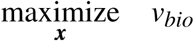

subject to:

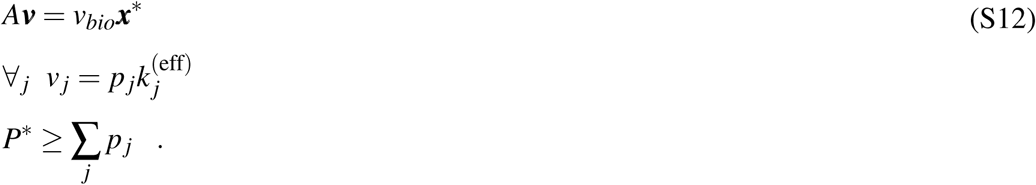

We relaxed the constraint (S3) on total protein into an inequality constraint, so that Eq. (S12) describes a protein-constrained FBA problem for the active stoichiometric matrix. This is precisely the type of constrained flux balance problem analyzed in Refs.^42, 43^, which prove that the solutions ***v***^opt^ to the optimization problem defined by Eq. (S12) are elementary flux modes (EFMs).

In the optimal solution to the problem defined by Eq. (S12), the protein concentration constraint will be active, that is, *P*\* = Σ_*j*_ *p*_*j*_; if not, the biomass production rate *v*_*bio*_ could be increased by multiplying the vector of protein concentrations ***p*** with a constant > 1 (as 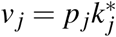 or all *j*). Thus, the optimization problem described by Eq. (S12) is the same as that described by Eq. (S11), except for a reduction in the dimension of the search space due to the fixed concentrations ***x***\* (Note that the cellular capacity constraint (S4) is trivially respected in Eq. (S12) and can be ignored). Accordingly, the flux distribution ***v***\* that maximizes the balanced growth rate *µ* in Eq. (S11) also maximizes the biomass production rate *v*_*bio*_ of the protein-constrained FBA problem in Eq. (S12); it is hence a representative of an EFM of the active stoichiometric matrix *A* with biomass ***x***\*,^42, 43^ and thus ***v***\* is an EGS according to Def. 3.

In parallel work to that presented here, de Groot *et al.* have shown that optimal solutions to balanced growth problems are elementary growth modes as defined in Ref.^18^, and that the active stoichiometric matrix of elementary growth modes has full rank^18^.

If instead of a single constraint on cellular capacity, multiple capacity constraints are imposed simultaneously (e.g., to describe separate constraints on different cellular compartments), then the solutions may in some cases correspond to positive linear combinations of EGSs^18, 45^, and the treatment below would have to be generalized. Multiple capacity constraints may play a role in the emergence of overflow metabolism in *E. coli*^46^, although overflow metabolism can also arise in balanced growth models with a single capacity constraint^5^.

In a cBGS with maximal growth rate for a given active stoichiometric *matrix A*, the cellular components will utilize the full cellular capacity *ρ* to saturate enzymes with their substrates. Thus, the constraint (S4) will be active, turning the inequality into an equality. The maximal balanced growth rate *µ*\* will thus be a function of the cellular capacity *ρ*. As a reference value for the total marginal net benefits of individual concentrations *x*_*i*_, we now define the marginal benefit of the cellular capacity *ρ* (constraint (S4)). This is the first definition that makes use of the capacity constraint.

##### Definition 5

(Marginal benefit of the cellular capacity). *In analogy to the marginal net benefits of cellular components, we define the* ***marginal benefit*** *of the cellular capacity as the fitness increase facilitated by a small increase in ρ*,

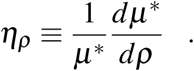

We can now relate *η*_*ρ*_ to the total marginal net benefits of all concentrations. To do this, we derive necessary conditions for any optimal balanced growth state at constant cellular capacity *ρ*, using the method of Lagrange multipliers. The Lagrange multipliers quantify the importance of the capacity constraint, Eq. (S4), and of the constraints for the dependent reactants, Eq. (S8), for the maximization of the objective function. The Lagrangian ℒ is a function of *P*, ***a***, and *ρ*.

##### Theorem 10

(Balance equation). *In a cBGS with maximal growth rate, the total marginal net benefit of each independent concentration x*_*i*_ ∈ {*P, b*_*β*_} *equals the marginal benefit of the cellular capacity ρ scaled by the capacity factor κ*_*i*_,

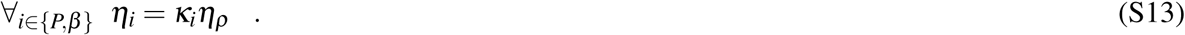

*Proof.* We use the method of Lagrange multipliers to derive necessary conditions for any optimal cBGS at constant cellular capacity *ρ*. Our objective function is given by Theorem 7, which expresses the growth rate *µ* as an explicit function of the concentrations ***x*** = [*P*, ***a***]^*T*^. The capacity constraint (S4) will be active at maximal growth rate, i.e., it becomes an equality. The capacity constraint can then be expressed as a function *g*_*ρ*_ that depends on *ρ* and on the concentrations,

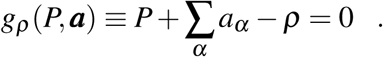

Finally, the constraints on each dependent reactant *γ* also only depend on *P*, ***a***, with the entries *D*_*γP*_ determining the composition of each *γ* in terms of *P*, and *D*_*γβ*_ determining the composition of *γ* in terms of *b*_*β*_,

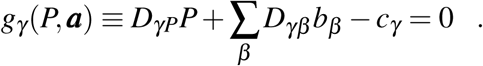

We now define a Lagrangian as the sum of the objective function *µ* and the constraints ***g*** scaled by Lagrange multipliers *λ*_*ρ*_, accounting for the capacity constraint (S4), and *λ*_*γ*_, accounting for the dependence of the dependent reactants *γ* ∈ {*γ*}, Eq. (S8):

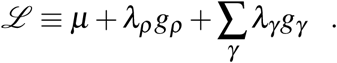

The first order necessary conditions for a constrained local maximum are that all partial derivatives of ℒ with respect to the variables *P, b*_*β*_, *c*_*γ*_ and to the Lagrange multipliers *λ*_*ρ*_, *λ*_*γ*_ are zero,

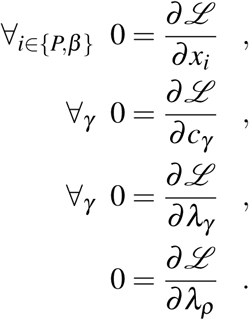

For the partial derivative with respect to an independent concentration *x*_*i*_ (*i* ∈ {*P, β*}), we have

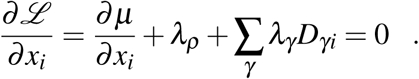

With Theorem (8), this results in

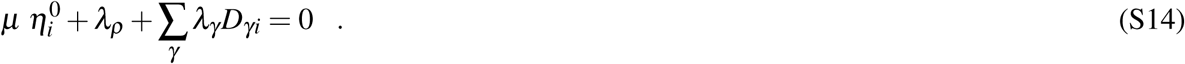

For the partial derivative relative to a dependent reactant *c*_*γ*_, we have

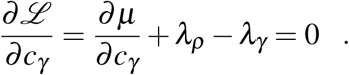

With Eq. (S9), we obtain

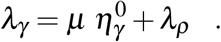

Substituting *λ*_*γ*_ from the last equation into Eq. (S14) gives (for *i* ∈ {*P, β*})

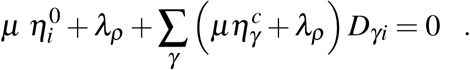

Rearranging results in

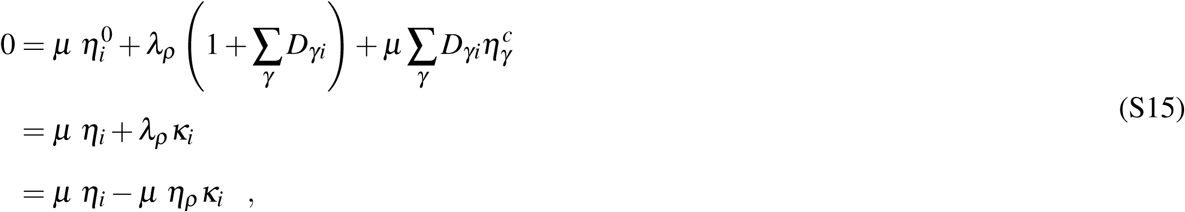

where we used *η*_*ρ*_ = −*λ*_*ρ*_ */µ*, which follows directly from the envelope theorem^47^. With *µ* > 0, we thus obtain the balance equation.

The optimal state is perfectly balanced: the total marginal net benefit of each independent cellular concentration *x*_*i*_ equals the marginal benefit of the cellular capacity, scaled by *κ*_*i*_ to account for its total utilization of cellular capacity. If *i* does not have any dependent reactants (∀_*γ*_ *D*_*γi*_ = 0), then the balance equation simplifies to 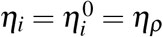 (Eq. (10)).

Theorem 10 states that if the dry weight density *ρ* would be allowed to increase by a small amount, such as 1 mg/l, then the marginal fitness gain that could be achieved by increasing protein concentration (plus dependent concentrations) by this amount is identical to that achieved by increasing the concentration of any reactant *β* (plus its dependent concentrations) by the same amount.

Instead of using Lagrange multipliers in the proof, one could express the total protein concentration *P* = *ρ* − Σ_*α*_ *a*_*α*_ (constraint (S4)) and the dependent reactant concentrations *c*_*γ*_ = *D*_*γP*_*P* + Σ_*β*_ *D*_*γβ*_ *b*_*β*_ (Eq. (S8)) in terms of *ρ* and of the independent reactant concentrations ***b***. Substituting the resulting expressions into the growth equation (Theorem 7) would result in an objective function that depends only on *ρ* and ***b***, and that is constrained only by the requirement of positive concentrations. While this would lead to the same balance equations as derived in the Lagrange multiplier framework, this formulation misses important insights that can be derived from the Lagrange multipliers themselves.

### B. Definition of relative fitness

In a situation where competition among cells is solely through differential intrinsic growth rates, absolute fitness is equal to growth rate: In a population of cells growing exponentially with growth rate *µ*, the selection coefficient for a variant with growth rate *µ* + *δ µ* is simply *δ µ*. ^1^ Population genetics models almost always employ relative fitness^48^, which we here define as a relative growth rate:

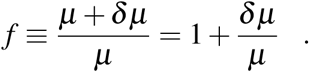

Thus, to quantify the effect on relative fitness of a small change of some parameter *x* by *δx*, we use

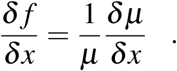

Note that population genetics models are frequently defined in terms of discrete generations. With generation time *T*_gen_ = ln 2*/µ*, the selection coefficient of the variant *per generation* is then^49^

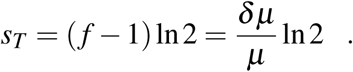

### C. Optimal ribosome protein fraction

Here we assume a very simple model for translation^26^. It accounts only for the elongation phase, where one catalyst (the ribosome plus bound mRNA, with concentration *R*) converts one substrate (the ternary complex, with concentration *a*_*T*_) into protein, following irreversible Michaelis-Menten kinetics:

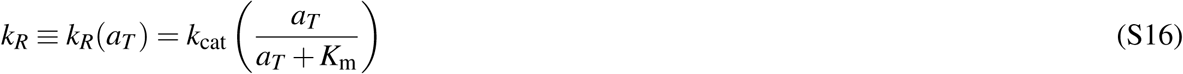

with constant maximal ribosome activity *k*_cat_ (in units of [time]^−1^) and Michaelis constant *K*_m_ (in units of [mass][volume]^−1^).

As further simplifications, we assume that the model has no dependent reactants (*A* = *B*) and that the ternary complex is not used in any other reaction. In this case, the same canceling of production costs as in the model depicted in Fig.S1A happens, and the balance of net benefits of ternary complex and total protein, *η*_*T*_ = *η*_*P*_ (Eq. (10)), simplifies to

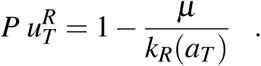

Substituting the partial derivative of irreversible Michaelis-Menten kinetics (Eq. (S16)), we obtain

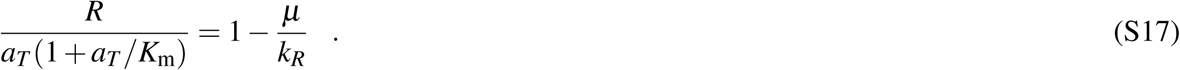

Rearranging Eq. (S16), we also see that the kinetics determine the concentration *a*_*T*_ uniquely in terms of *v*_*R*_, *R, K*_m_, and the ribosome’s turnover number *k*_cat_,

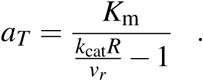

Substituting this into Eq. (S17) gives

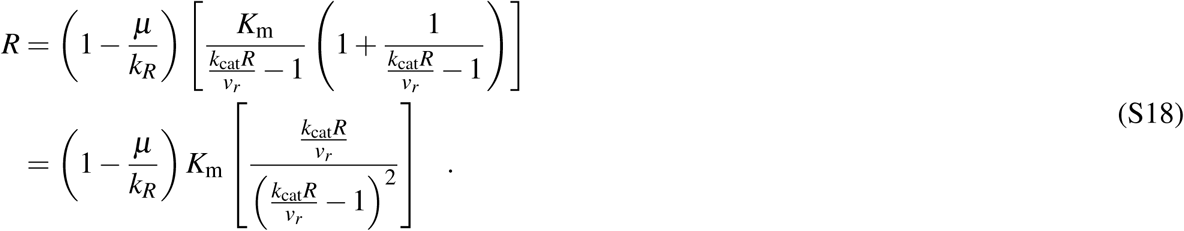

From the ribosome kinetics and mass conservation of proteins, we have

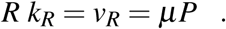

Thus, substituting *µ/k*_*R*_ = *R/P* and *v*_*R*_ = *µP* in Eq. (S18), we obtain

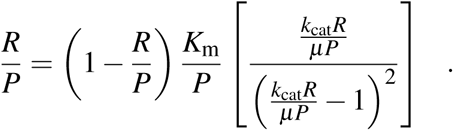

This is equivalent to a quadratic equation in *R/P*,

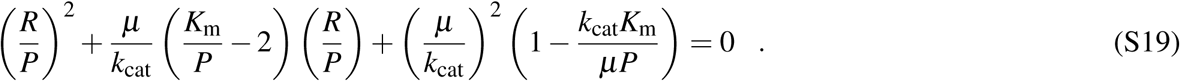

Its two solutions are

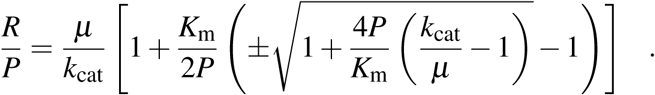

To see which of the two solutions is relevant, we rewrite this as

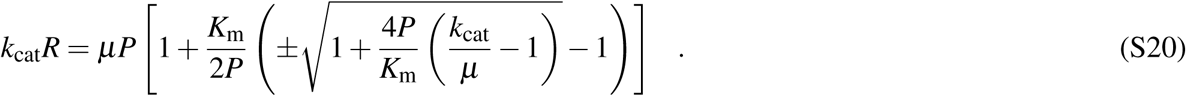

Because *k*_cat_*R* > *R k*_*R*_ = *v*_*R*_ = *µP*, the term in square brackets ([·]) in Eq. (S20) must be > 1. Only the positive root is compatible with this condition. Thus, the ratio *R/P* is uniquely determined by

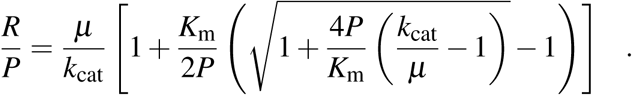

To estimate the actual ribosome protein fraction of total protein *ϕ*_*R*_, we need to scale the previous expression by the fraction *r*_*P*_ of ribosome which is protein, resulting in the final equation

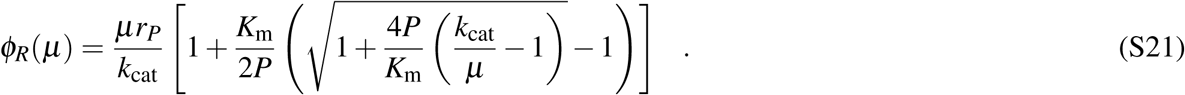

The same procedure can be used to find an equation for *ϕ*_*R*_ that ignores the production costs. Starting from Eq. (S18) without the production cost term *µ/k*_*R*_, we obtain

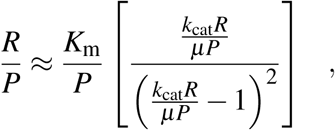

which results in a quadratic equation similar to Eq. (S19),

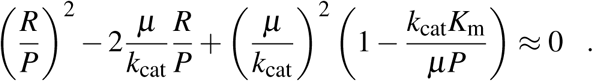

Solving for *R/P* gives

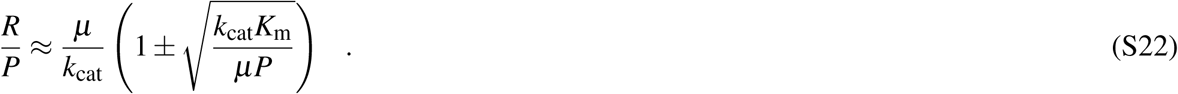

Again because *Rk*_cat_ > *µP*, the term in parentheses (·) in Eq. (S22) must be > 1, and again only the positive root is compatible with this condition. Thus, the ribosome protein fraction is uniquely determined in this approximation by

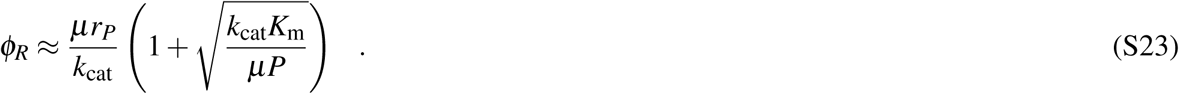

We compared the predictions of the ribosome fraction of total protein, *ϕ*_*R*_ = *R/P*, to quantitative proteomics data obtained by Schmidt *et al.*^32^ (scaled by a factor of 0.67, as suggested by the authors, to account for a systematic error in the cell size measurements^50^). To obtain molar ribosome concentration, we calculated the median over all reported concentrations of ribosomal proteins. The concentration of ternary complexes was assumed to be identical to the concentration of their protein component, the elongation factor Tu. Molar concentrations of the ribosome and (total) ternary complexes were converted to mass concentrations by multiplying with molar masses derived from the amino acid sequences (for the protein parts) and nucleotide sequences (for the RNA parts). For this, we assumed that each ribosome contained one copy of each of its constituents, with the exception of four copies of RplL^51^. To calculate the mass fraction of total protein occupied by ribosomes, we multiplied ribosome mass concentrations with the mass fraction of ribosomes that is protein *(r*_*P*_ = 0.58^32^), and divided the result by the total protein mass concentration *P* = 127.4 g/l in *E. coli*, assumed to be constant across growth conditions^32^.

The concentration of actively translating ribosomes was determined based on total ribosome concentration and the fraction of active ribosome at different growth rates. The latter was estimated by fitting a smooth saturation function *s*(*µ*) = *µ/*(*µ* + *z*) over the fractions of active ribosomes estimated in Ref.^33^, with the best-fitting parameter *z* = 0.124*/h*.

We set the Michaelis constant of the ribosome to *K*_m_ = 3 × 10^−6^ mol/l, based on the diffusion limit for ternary complexes calculated in Ref.^26^. We set the ribosome’s turnover number to *k*_cat_ = 22 AA/s, the highest elongation rate observed experimentally in Ref.^52^. As we do not distinguish between different ternary complexes and the ribosome only accepts one of the 40 different ternary complex types at any given time, *K*_m_ was multiplied by 40.^26^ For consistency of the units with the mass concentration units used throughout our paper, the kinetic parameters had to be converted from molar to mass concentrations. The mean weight (± SD) of amino acids across all conditions assayed in Ref.^32^ was 132.60 ± 0.09 Da; the ribosome molecular weight is 2, 306, 967 Da; and the mean weight of ternary complexes is 69, 167 ± 1, 351 g/mol. With these numbers, we obtain *k*_cat_ = 22 AA/s × (132.60 Da/AA)/(2,306,967 Da) ×3, 600s/1h = 4.55/h, and *K*_m_ = 40 × 3 × 10^−6^mol/l ×69, 167 g/mol = 8.30 g/l.

### D. Minimal whole-cell model and the dependence of maximal growth rate on cellular water content

Cayley *et al.*^29, 35^ showed that the internal water content of *E. coli* cells increases when these are grown in environments with reduced osmolarity. This effect corresponds to a decrease of cellular dry weight per volume, *ρ*, by *δρ. η*_*ρ*_ quantifies the associated reduction in relative fitness, *δ f* = *δ µ*\**/µ*\* = *η*_*ρ*_ *δρ*, with *µ*\* the maximal growth rate (Def. 5). The relative change in the maximal growth rate per relative change in *ρ* is then

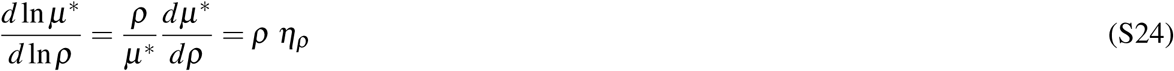

From Eq. (S13), we know that *η*_*P*_ = *κ*_*P*_*η*_*ρ*_; if there are no dependent reactants for *P* (i.e., ∀_*γ*_ *D*_*γP*_ = 0), this simplifies to

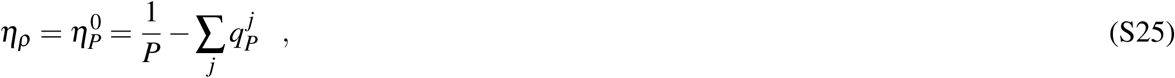

and thus

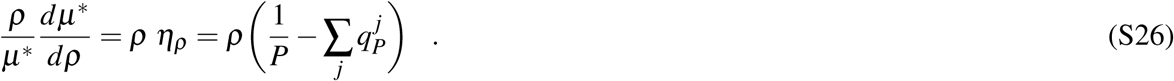

The mass fraction of total protein in cell dry weight *P/ρ* ≈ 0.54 has been shown to be approximately constant across growth conditions supporting intermediate to high growth rates^29^. To estimate the total protein production cost 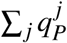, we consider the simplest possible whole-cell model, comprising only a transport reaction and the ribosome reaction (Fig. S2).

The active stoichiometric matrix *A* of this model and its inverse I = *A*^−1^ are, written here with row and column labels,

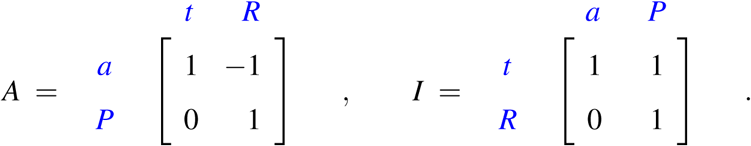

The capacity is determined only by its two components,

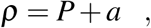

where

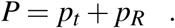

From the inverse *I* and Eq. (4), we obtain

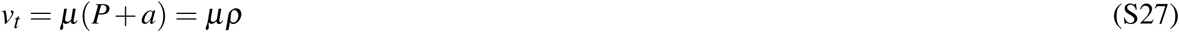

and

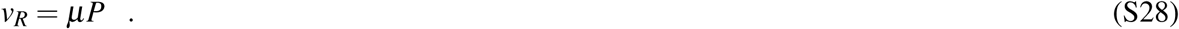

From the inverse *I* and Eq. (3), we get

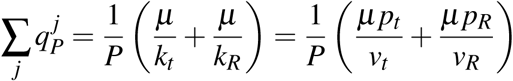

Combining this with Eq. (S27) and (S28) and with *ϕ*_*R*_ = *p*_*R*_*/P* and *ϕ*_*t*_ = *p*_*t*_ */P* = 1 −*ϕ*_*R*_, we obtain

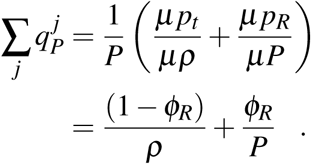

Combining this equation with Eq. (S26), we obtain

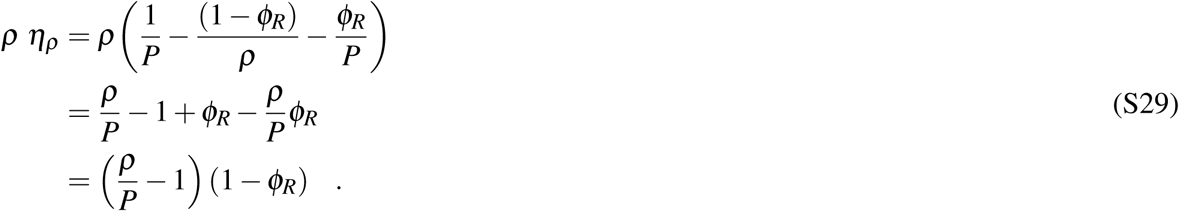

From Eq. (S21), we estimate the mass fraction of ribosomal proteins in total protein *ϕ*_*R*_ at *µ* = 1.0*/h* (growth rate in the reference growth condition of osmolarity Osm = 0.28 in Ref.^35^) as *ϕ*_*R*_ = 0.19. Substituting this value into Eq. (S29) together with *P/ρ* = 0.54, we estimate the relative change in the maximal growth rate per relative change in *ρ* as

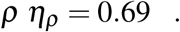

Cayley *et al.*^35^ report cell growth at reduced osmolarities, summarized in Table S1. The cell free water content 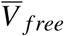 in Table S1 is calculated from the total cell water 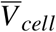 minus the observed constant bound water 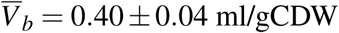.^29^ Errors are estimated standard deviations based on error propagation among normally distributed random variables.

Fig. S3 plots the natural logarithms of *µ* and *ρ*. Linear regression over the three available data points results in an estimated slope of 0.66, close close to our estimate of 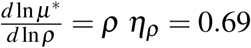.

### E. An outline of possible extensions of GBA

In our development of GBA, we make several simplifying assumptions. Here, we outline some possible generalizations.

#### All proteins contribute to growth by acting as catalysts or transporters

This assumption can simply be removed by adding a sector of non-growth related proteins^25, 27^ with concentration *Q* to the r.h.s. of Eq. (2).

#### Proteins are not used as reactants

To use protein *j* as a reactant in reaction *j*′, it will need an extra row in *A*, and its concentration *p*_*j*_ has to enter the concentration vector 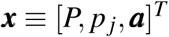 and the kinetic function *k*_*j*′_ (*p*_*j*_, ***a***). This does not affect Eq. (4). However, if *p*_*j*_ appears on the right hand side of Eq. (5), this equation will have to be solved for *p*_*j*_ before it is possible to proceed to a generalization of the growth equation.

#### All catalysts are proteins

We can add different catalytic RNA species as cellular components. Additionally, we may introduce reactions that combine proteins and RNA into molecular machines such as the ribosome.

#### A 1-to-1 correspondence between proteins and reactions

Spontaneous reactions that proceed without a catalyst have to be included in the active stoichiometric matrix *A* (so that *I* accounts for their dilution). They will need a kinetic function that relates their flux to the substrate concentrations (e.g., through mass action kinetics). However, they will not contribute to the protein sum (Eq. (2)) and hence will not directly contribute to the growth equation (6). Because in this case the flux cannot be adjusted by varying the concentration of a catalyst, only concentration vectors are feasible for which the flux through this reaction is identical when calculated based on mass conservation (through *I*) and on kinetics. This will reduce the dimensionality of the solution space.

In the case of isoenzymes, where both protein *j* and protein *j*′ catalyze the same reaction, the optimal solution will always use the one with the more favourable kinetics at the given concentrations (e.g., protein *j* if *k*_*j*_(***a***) > *k*_*j*′_ (***a***) > 0).

For protein complexes, where proteins *j* and *j*′ have to bind to each other before they can act as a catalyst, we can either ignore the individual proteins and include the protein complex as a cellular component in the model, or add a reaction that describes the complex formation.

Finally, if one protein (or protein complex) catalyzes reactions *j* and *j*′, the substrates (and possibly products) of reaction *j*′ will enter the kinetic function *k*_*j*_(***a***). The fluxes through both reactions are proportional to the protein concentration *p*. Hence, *p* = *v*_*j*_*/k*_*j*_(***a***) = *v*_*j*′_*/k*_*j*′_ (***a***), providing an additional constraint for the fluxes *v*_*j*_, *v*_*j*′_. As the fluxes are unique given the concentration vector ***x*** = [*P*, ***a***]^*T*^, again not all concentration vectors ***x*** will be compatible with this condition, reducing the dimensions of the solution space of balanced growth.

#### Optimizing only growth

An extension of GBA can be formulated for non-growing cells (or cellular subsystems) that are instead optimized for the production of specific molecules, as is the case for many cell types in multicellular organisms. The dilution term *µ****x*** in Eq. (1) would be replaced by a vector ***d***(***x***) that quantifies the degradation of proteins and other molecules (with entries *d*_*i*_ = *z*_*i*_*x*_*i*_ and constant degradation rate *z*_*i*_); an additional “output vector” ***o*** would represent the desired cellular production, with rate *v*_*o*_:

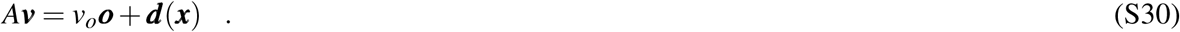

The kinetics are still a function of ***x***, and we can proceed with the analysis following the same steps as for Eq. (1) to calculate ***v, p***, and *v*_*o*_.

### F. Growth Control Analysis (GCA)

Here, we briefly explore the connection between GBA and some central concepts of *metabolic control analysis* (MCA)^16^. The results below that involve elasticity and control coefficients largely restate previous insights^22^, 23 in the framework of GBA. First, we rephrase the balance equation in terms of control theory.

We define the (scaled) *growth control coefficients* (GCC) as the *total* relative change in the growth rate due to a small change in the concentration *x*_*i*_, *accounting for the capacity constraint*. The growth rate change is caused by two effects: the net fitness benefit of increasing *x*_*i*_ without considering the capacity constraint, captured by the marginal net benefits *η*_*i*_; and the fitness cost of reducing the cellular capacity *ρ* available for all other concentrations, captured by −*κ*_*i*_*η*_*ρ*_. The GCC is then simply the sum of these two,

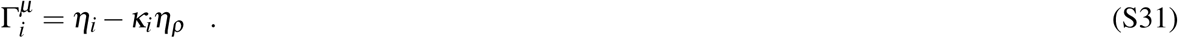

From the balance equation, we have 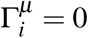 at optimal growth. Growth control coefficients have been defined before for changes in growth rate due to small changes in the concentration of individual proteins, in the context of noise propagation in a model of gene expression and cellular growth^21^.

We now examine the the relationship of the variables defined in GBA to the coefficients considered in MCA. The *elasticity coefficient* 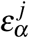 in MCA is defined as the change in the reaction rate *j* when varying the the substrate concentration *a*_*α*_ while keeping the enzyme (catalyzing protein) concentration fixed^16^. The *normalized* elasticity coefficient is thus directly related to the marginal kinetic benefit 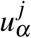,

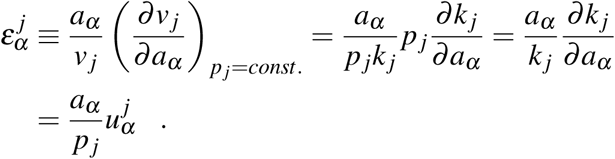

Control coefficients have been defined in MCA as the change in a *response variable y* due to a change in a *state variable x*, where each *y* = *y*(***x, π***) is a function of the state variables ***x*** and the *system parameters* ***π***^16^. In the GBA framework, the growth rate *µ*, the fluxes ***v***, the protein concentrations ***p***, and the dependent concentrations ***c*** are all functions of the concentrations ***x*** = (*P*, ***b***), the active matrix *A*, and the kinetic parameters in ***k***, Thus, *µ*, ***v, p***, and ***c*** can be seen as response variables, while the concentrations in ***x*** are state variables. In contrast to MCA, the GBA framework provides explicit functions for all response variables, and thus control coefficients can be calculated easily. The control of *µ* by the concentrations *x*_*i*_ is given by the growth control coefficient 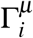 in Eq. (S31), while the control of dependent concentrations *c*_*γ*_ is directly determined by the dependence matrix *D*.

We next examine the control of fluxes *v*_*j*_ and protein concentrations *p*_*j*_. The (scaled) *flux control coefficient* (FCC) 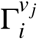 is the relative change in *v*_*j*_ due to a small change in *x*_*i*_ (at fixed concentrations *x*_*i*′_ for *i*′ ≠*i*),

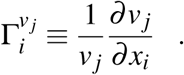

From Eq.(4), we can calculate *∂v*_*j*_*/∂x*_*i*_, giving

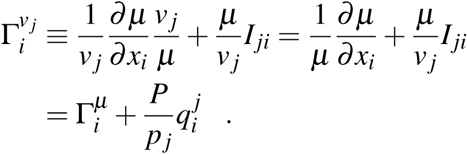

At optimal growth, 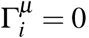, so

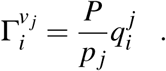

Thus, at optimal growth, the flux control coefficient is simply the marginal production cost incurred via reaction *j*, divided by the protein fraction of the catalyzing protein.

The (scaled) *protein control coefficient* (PCC) 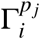 is the change in the protein fraction of protein *j, p*_*j*_*/P*, due to a small change in the concentration *x*_*i*_ (at fixed concentrations *x*_*i*′_ for *i*′ *≠ i*),

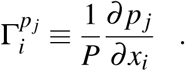

From the kinetic constraint (S2),

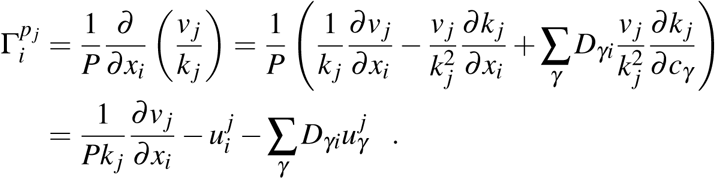

Again calculating *∂v*_*j*_*/∂x*_*i*_ from Eq. (4), we obtain

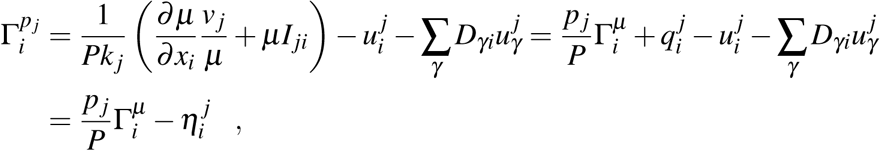

where we defined 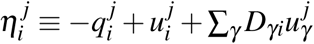 as the contribution of reaction (or protein) *j* to the marginal net benefit *η*_*i*_. Summing over *j*, we obtain

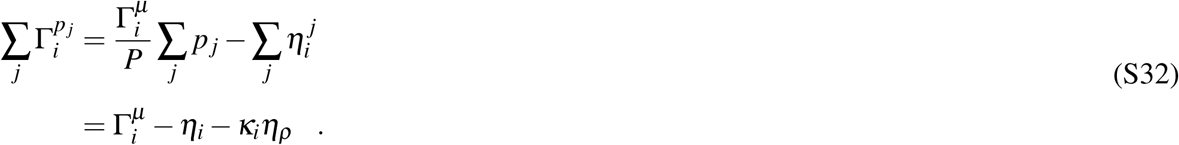

Without a capacity constraint, *η*_*ρ*_ = 0, and

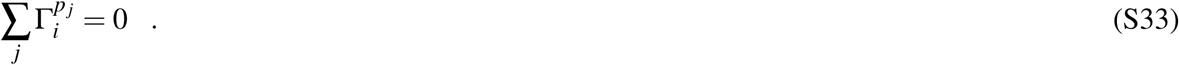

Equations (S32) and (S33) can be seen as *summation theorems* that relate the GCC 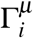 with the control coefficients of MCA, in a similar fashion as in^21^.

At optimal growth,

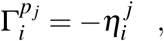

and

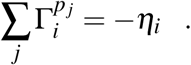

Typically, reactants participate in only a small fraction of reactions, so for most combinations 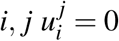 and 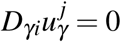; the PCC at optimal growth is then just the marginal production cost,

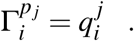

### G. Choice of basis and relationship between capacity and dependence constraints

Not every reactant can be considered dependent: a reactant for which the corresponding row in the active matrix *A* is linearly independent of all other rows will always be in the basis (equivalently, a reactant that has zero entries in all vectors in a basis for the left null space of *A* cannot be a dependent reactant).

It is possible for some models that there is one or more choices of basis such that its corresponding dependence matrix has for some *i* ∈ {*P, β*}

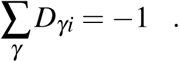

In these cases, any marginal change in the mass concentration of component *i* will cause the exact opposite change in the total mass concentration of its dependent reactants *γ*. When this is combined with the capacity constraint as defined in Eq. (9), these changes in concentrations result in a perfect cancellation in the capacity utilized by *i* and its dependent reactants, and thus a zero net change in capacity for any change in the concentration *i* (i.e., *κ*_*i*_ = 0, Def. 4). For this reason, the marginal net benefit of *i* is simply *η*_*i*_ = 0 (Eq. (10)).

Such a perfect cancellation is highly unlikely if we use a more realistic description of the capacity constraint, where different cellular components *i* have different specific capacity utilizations *σ*_*i*_; e.g., if we assume that the capacity constraint limits the total volume occupied by cellular components, then *σ*_*i*_ gives the volume per mass of component *i*. In this case, the capacity constraint Eq. (9) is replaced by a constraint on the volume of cellular dry mass per volume of cell water, *v*:

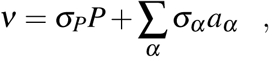

where *σ*_*P*_ is the specific capacity of proteins (almost constant for different proteins^53^) and *σ*_*α*_ is the specific capacity of reactant *α*, which depends on its chemical properties such as hydrophobicity and charge^54^.

## Supplementary Figures

**Figure S1.**
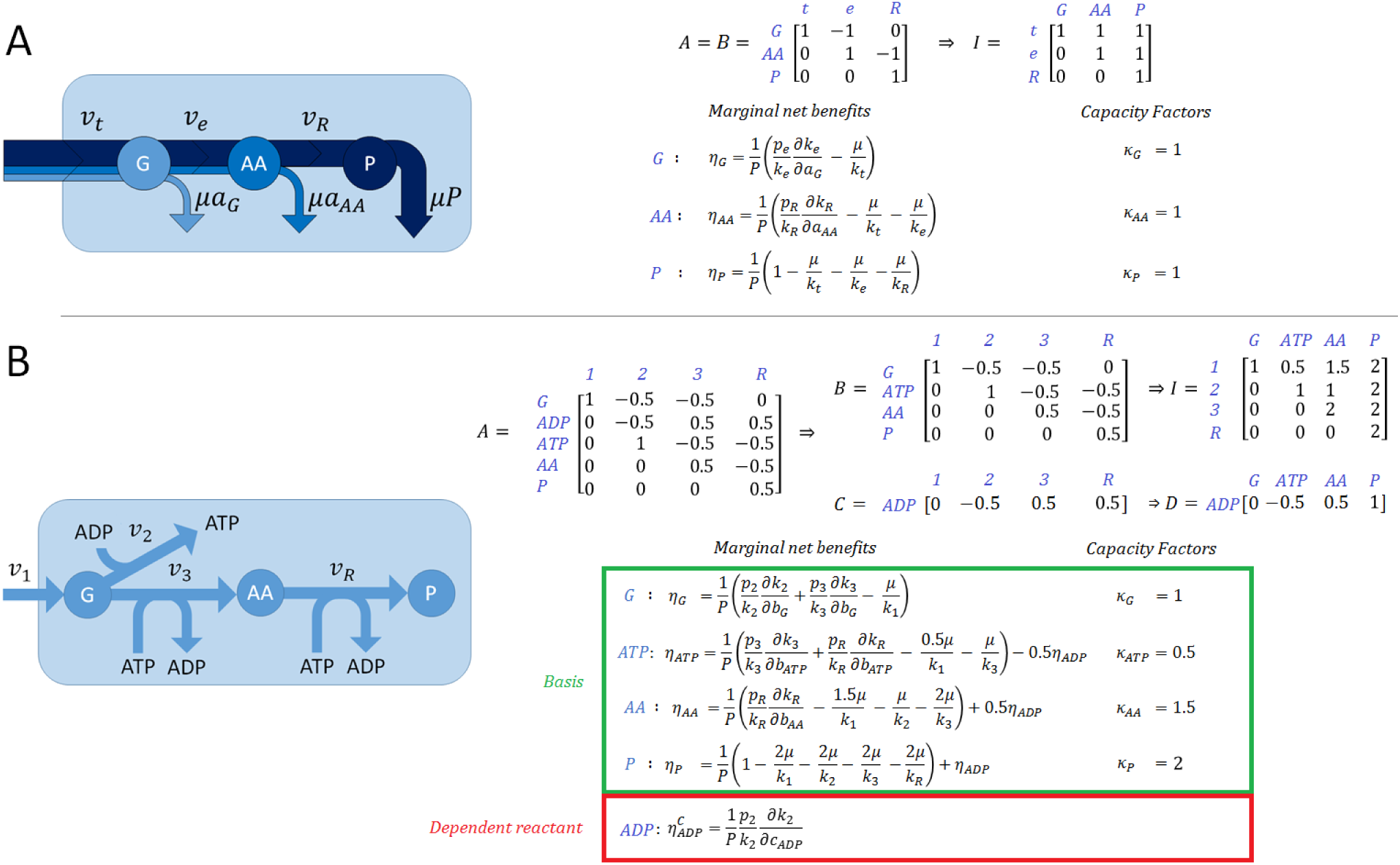
Examples of balanced growth models and their mathematical description, derived from the active matrix *A* and the kinetic functions *k*_*j*_(***a***): basis matrix *B*, investment matrix *I* = *B*^−1^, closure matrix *C*, dependence matrix *D* = *CI*, marginal net benefits *η*_*i*_, and capacity factors *κ*_*i*_. (A) A model with a simple linear network of irreversible reactions, connecting a single transporter to the final production of proteins; linear networks never have dependent reactants, as the number of reactions equals the number of components (*n* = *m* + 1). Colors indicate the fraction of flux that is eventually diverted into the dilution of each downstream component. (B) A more elaborate, nonlinear model of irreversible reactions that includes cofactors and a dependent reactant (ADP).

**Figure S2.**
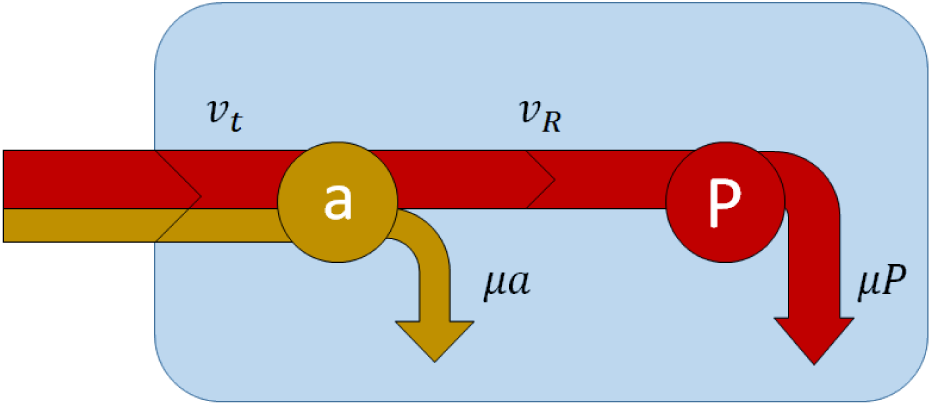
Minimal whole-cell model, comprising a transport reaction (with rate *v*_*t*_) and the ribosome reaction (with rate *v*_*R*_).

**Figure S3.**
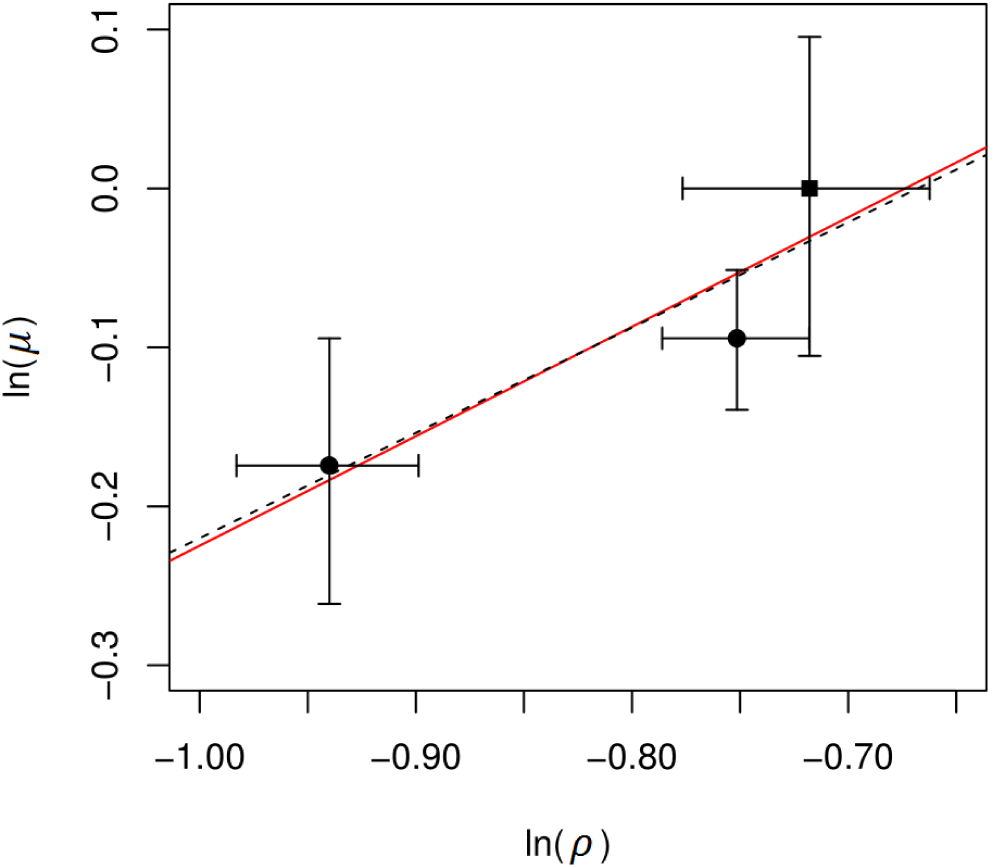
Dependence of the growth rate (ln *µ*) on dry weight per free water volume (ln *ρ*) in *E. coli* grown at different external osmolarities^35^. The square (▪) indicates the normal environmental conditions, which correspond to the maximal growth rate; dots (•) indicate growth at lower osmolarities. The dotted line indicates the linear regression with slope = 0.66. The red line indicates the predicted slope = 0.69, drawn through the center of gravity of the 3 data points. Error bars are based on the reported experimental standard deviations.

**Figure S4.**
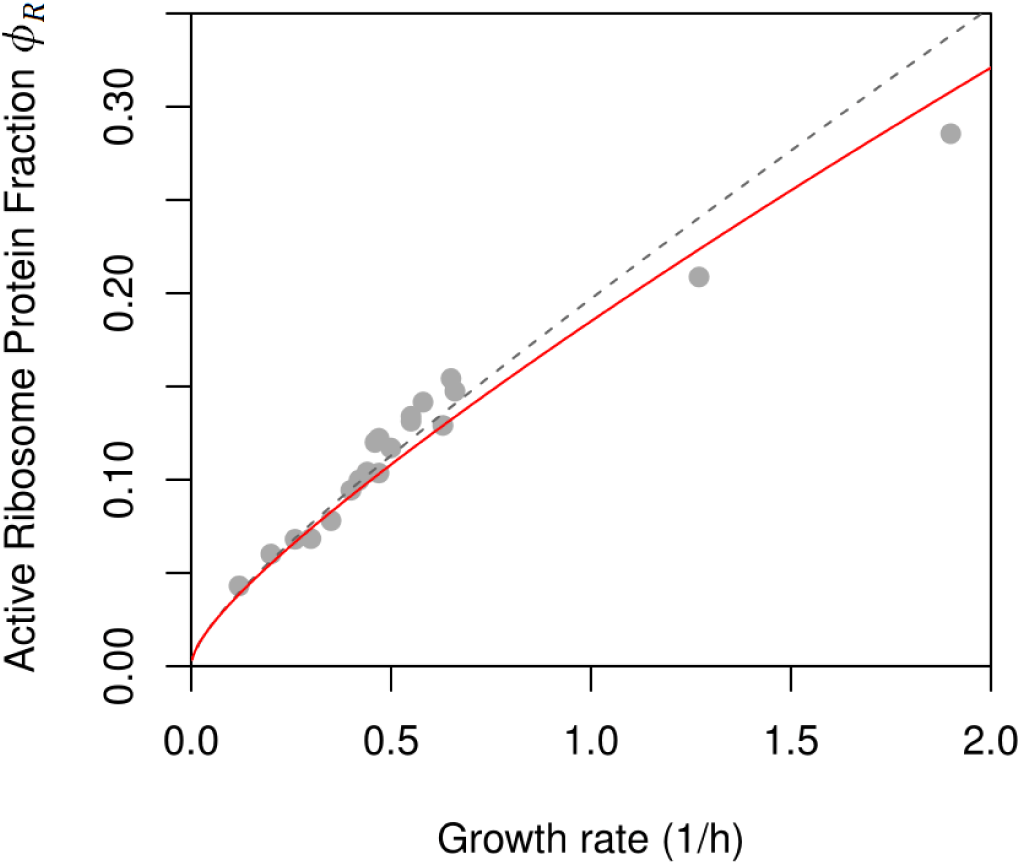
An approximation (dashed grey line, no free parameters; Eq. (S23)) that ignores the dilution of intermediates and hence production costs 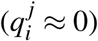 results in good predictions of experimentally observed *E. coli* active ribosome protein fraction^32, 33^ at low to intermediate growth rates (see also Ref.^34^). For comparison, we also show the full GBA prediction (red line, identical to Fig. 2).

## Supplementary Tables

**Table S1.** Experimental data from Cayley *et al.*^35^, including cellular free water content 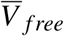 and growth rate *µ* across different external osmolarities, together with the respective cellular dry weight per cellular free water volume, calculated as 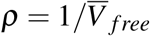.

| Osmolarity (Osm) | $\bar{V}_{free}$ (ml/gCDW) | $\rho$ (gCDW/ml) | $\mu$ (1/h) |
| --- | --- | --- | --- |
| 0.03 | $2.56 \pm 0.10$ | $0.39 \pm 0.02$ | $0.84 \pm 0.07$ |
| 0.10 | $2.12 \pm 0.08$ | $0.47 \pm 0.02$ | $0.91 \pm 0.04$ |
| 0.28 | $2.05 \pm 0.11$ | $0.49 \pm 0.03$ | $1.00 \pm 0.10$ |

**Table S2.** Symbols and definitions. For simplicity of notation, we also use *P* as an index for total protein, and *ρ* as an index for cellular dry weight per volume.

| Symbol | Definition (units) |
| --- | --- |
| $A$ | Active matrix [mass fraction] |
| $B$ | Basis matrix [mass fraction] |
| $C$ | Closure matrix [mass fraction] |
| $D$ | Dependence matrix |
| $I$ | Investment matrix |
| $v$ | Reaction rate [mass][volume]<sup>-1</sup>[time]<sup>-1</sup> |
| $\mu$ | Growth rate [time]<sup>-1</sup> |
| $P$ | Total protein concentration [mass][volume]<sup>-1</sup> |
| $a$ | Reactant concentration [mass][volume]<sup>-1</sup> |
| $b$ | Basis reactant concentration [mass][volume]<sup>-1</sup> |
| $c$ | Dependent reactant concentration [mass][volume]<sup>-1</sup> |
| $\alpha$ | Reactant index |
| $\beta$ | Basis reactant index |
| $\gamma$ | Dependent reactant index |
| $j$ | Reaction index |
| $i$ | Protein and active reactant index ($\{P, \beta\}$) |
| $k$ | Kinetic function (in units of $k_{\text{cat}}$) |
| $k_{\text{cat}}$ | Turnover number [time]<sup>-1</sup> |
| $K_m$ | Michaelis constant [mass][volume]<sup>-1</sup> |
| $\rho$ | Cellular dry weight per volume [mass][volume]<sup>-1</sup> |
| $f$ | Fitness |
| $\eta_i^0$ | Direct marginal net benefit of $i$ [volume][mass]<sup>-1</sup> |
| $\eta_i$ | Marginal net benefit of $i$ [volume][mass]<sup>-1</sup> |
| $\eta_\rho$ | Marginal benefit of of the cellular capacity [volume][mass]<sup>-1</sup> |
| $\eta_\gamma^c$ | Marginal net benefit of $\gamma$ [volume][mass]<sup>-1</sup> |
| $q_i^j$ | Marginal production cost of $i$ relative to $j$ [volume][mass]<sup>-1</sup> |
| $u_\beta^j$ | Marginal kinetic benefit of $\beta$ relative to $j$ [volume][mass]<sup>-1</sup> |
| $u_\gamma^j$ | Marginal kinetic benefit of $\gamma$ relative to $j$ [volume][mass]<sup>-1</sup> |
| $\kappa$ | Capacity factor |
| $\mathcal{L}$ | Lagrangian |
| $\lambda_\rho$ | Lagrange multiplier of the capacity constraint |

